# Single Cell Atlas of Human Dura Reveals Cellular Meningeal Landscape and Insights into Meningioma Immune Response

**DOI:** 10.1101/2021.08.03.454066

**Authors:** Anthony Z. Wang, Jay A. Bowman-Kirigin, Rupen Desai, Pujan R. Patel, Bhuvic Patel, Saad M. Khan, Diane Bender, M. Caleb Marlin, Jingxian Liu, Joshua W. Osbun, Eric C. Leuthardt, Michael R. Chicoine, Ralph G. Dacey, Gregory J. Zipfel, Albert H. Kim, Allegra A. Petti, Gavin P. Dunn

**Affiliations:** Department of Neurological Surgery, Washington University School of Medicine, St. Louis, Missouri; Department of Pathology and Immunology, Washington University School of Medicine, St. Louis, Missouri; Andrew M. and Jane M. Bursky Center for Human Immunology and Immunotherapy Programs, Washington University School of Medicine, St. Louis, Missouri; Brain Tumor Center, Washington University School of Medicine/Siteman Cancer Center; Washington University School of Medicine, St. Louis, Missouri; Arthritis & Clinical Immunology Human Phenotyping Core, Oklahoma Medical Research Foundation, Oklahoma City, Oklahoma; Department of Genetics, Washington University School of Medicine, St. Louis, Missouri; McDonnell Genome Institute, Washington University School of Medicine, St. Louis, Missouri; Department of Medicine, Washington University School of Medicine, St. Louis, Missouri

## Abstract

Recent investigation of the meninges, specifically the dura layer, has highlighted its importance in CNS immune surveillance beyond a purely structural role. However, most of our understanding of the meninges stems from the use of pre-clinical models rather than human samples. In this study, we use single cell RNA-sequencing to perform the first characterization of both non-tumor-associated human dura and meningioma samples. First, we reveal a complex immune microenvironment in human dura that is transcriptionally distinct from that of meningioma. In addition, through T cell receptor sequencing, we show significant TCR overlap between matched dura and meningioma samples. We also identify a functionally heterogeneous population of non-immune cell types and report copy-number variant heterogeneity within our meningioma samples. Our comprehensive investigation of both the immune and non-immune cell landscapes of human dura and meningioma at a single cell resolution provide new insight into previously uncharacterized roles of human dura.

## Introduction

The central nervous system (CNS) in vertebrates is encased by three layers of tissue that together comprise the meninges(*1*). The outermost layer of tissue is the dura mater, the arachnoid mater lies one layer deeper, and the innermost layer is the pia mater, that adheres to the brain surface. The dura layer performs important structural roles in the CNS(*2*). Specifically, this layer protects the underlying brain and spinal cord, harbors the large vascular sinus and lymphatic vessels through which venous and lymphatic drainage of the brain traverses, and creates intracranial compartments that divide the cerebral hemispheres and separate them from the cerebellum of the posterior fossa. Combined, the dura and arachnoid mater form a water-tight seal to contain cerebrospinal fluid which originates from the choroid plexus and bathes the brain before exiting through arachnoid granulations. Thus, the dura is critical in establishing the anatomic compartments of the brain as well as other protective roles.

Beyond its structural roles, the meninges consist of cells which also perform critical functional roles in the CNS. The embryonic meninges influences development of the skull, neuronal migration and anatomic positioning, neurogenesis and blood vessel development, and the establishment of basement membranes of the pia and the glia limitans [reviewed in(*1, 3*)]. Recently, there has been a growing appreciation that the dura also harbors vital immunologic functions, in addition to its role as a physical barrier in the innate immune response of the brain, thereby supporting the view that the meninges represent a dynamic immune microenvironment involved in organizing CNS immune responses. First, several studies in mice have shown that the dura harbors a range of immune cell types including macrophages, monocytes, dendritic cells (DCs), T and B cells(*4*–*6*). Second, meningeal immunity is critical to immune responses against stroke, traumatic brain injury, infection, and cancer(*7*–*10*). Finally, recent identification of lymphatic channels in the dura has illuminated new mechanisms by which the CNS intersects with systemic immunity(*11, 12*). Several examples have demonstrated how modulating various functions of the dura could alter the immune response. For example, investigators showed that the CNS anti-tumor immune response could be attenuated by ligation of cervical lymphatics originating from the dura(*3*) and conversely, that CNS anti-tumor immune responses could be enhanced by the induction of dural lymphangiogenesis(*13*). Furthermore, clinicians have explored endovascular embolization of the middle meningeal artery—which perfuses the dura—for the treatment of chronic subdural hematomas(*14*). Thus, clarifying the cellular composition of dura may enable a better understanding of this tissue site, with important translational implications.

Because much of our understanding of meningeal biology stems almost entirely from preclinical models, we focused our work on characterizing meningeal composition in patients undergoing surgery for the resection of intracranial meningiomas. Meningiomas are common, typically benign, tumors originating from within the meninges and treated by surgical resection of the meningioma and nearby surrounding margin of dura, some of which is not associated with the tumor as determined by the surgeon(*15*). Herein, we report the first characterization of human dura using single cell RNA sequencing (scRNA-seq) by profiling the surrounding non-tumor-associated dura and a subset of matched meningiomas from patients undergoing surgical resection. We show that human dura consists of a diverse population of immune, endothelial, and mesenchymal cell types through our scRNA-seq analysis. We supplemented these observations with imaging mass cytometry (IMC), which allowed us to investigate the spatial relationships among these cell types. Moreover, from the scRNA-seq data, we observed cellular heterogeneity and functional diversity within each cell population characterized. In patient-matched dura and meningioma tumors, we observed that immune cell states were distinct within each tissue. Additionally, using single cell TCR sequencing, we show that dura which is tumor-adjacent, but not tumor-attached, harbors clonotypic T cell diversity and shares T cell clonotypes with adjacent meningioma tumor tissue. Finally, we provide the first evidence of copy number heterogeneity in primary meningioma tumor samples at the single cell level. Together, these findings provide further support that the dura is a dynamic anatomic tissue site and suggest cellular pathways by which the immune response to meningiomas evolve.

## Results

### The dura consists of a diverse landscape of both immune-related and non-immune-related cell types

To better understand the cellular composition of human dura, we performed scRNA-seq on samples of human dura and a subset of matched meningiomas derived from patients undergoing craniotomy for resection of intracranial meningiomas, which arise from the dura and thus are anatomically attached to this meningeal layer (Table S1). In surgical resection of convexity meningiomas, an adjacent region of uninvolved dura, as defined by the surgeon, is normally resected to ensure maximal tumor resection and reduce the risk of recurrence. This uninvolved, non-tumor-bearing dura was subsequently harvested and used in our analyses. Seven dura samples and four matched meningioma samples were disaggregated and analyzed using scRNA-seq (Fig. 1A). We first characterized the non-tumor-associated dura samples, performing unsupervised clustering and uniform manifold approximation and projection (UMAP) analysis on 22,460 cells (Fig. 1B). Cells were initially classified into 3 cell populations using common markers for endothelial cells (*PECAM1, CDH5, KDR*), mesenchymal cells (*COL1A1, COL1A2, LUM, DCN, ACTA2, RGS5*), and immune cells (*PTPRC, CD3E, SPI1, CD14*) (Fig. 1C and Table S2). The majority of cells were immune cells (10,423 cells), followed by endothelial cells (6,283 cells) and mesenchymal cells (5,754 cells). We identified each general cell type in all seven dura samples, indicating the absence of patient-specific clustering of cells (Fig. 1D). These data demonstrate that the dura harbors a diverse cell population of both immune and non-immune derived cell types.

**Figure 1:**
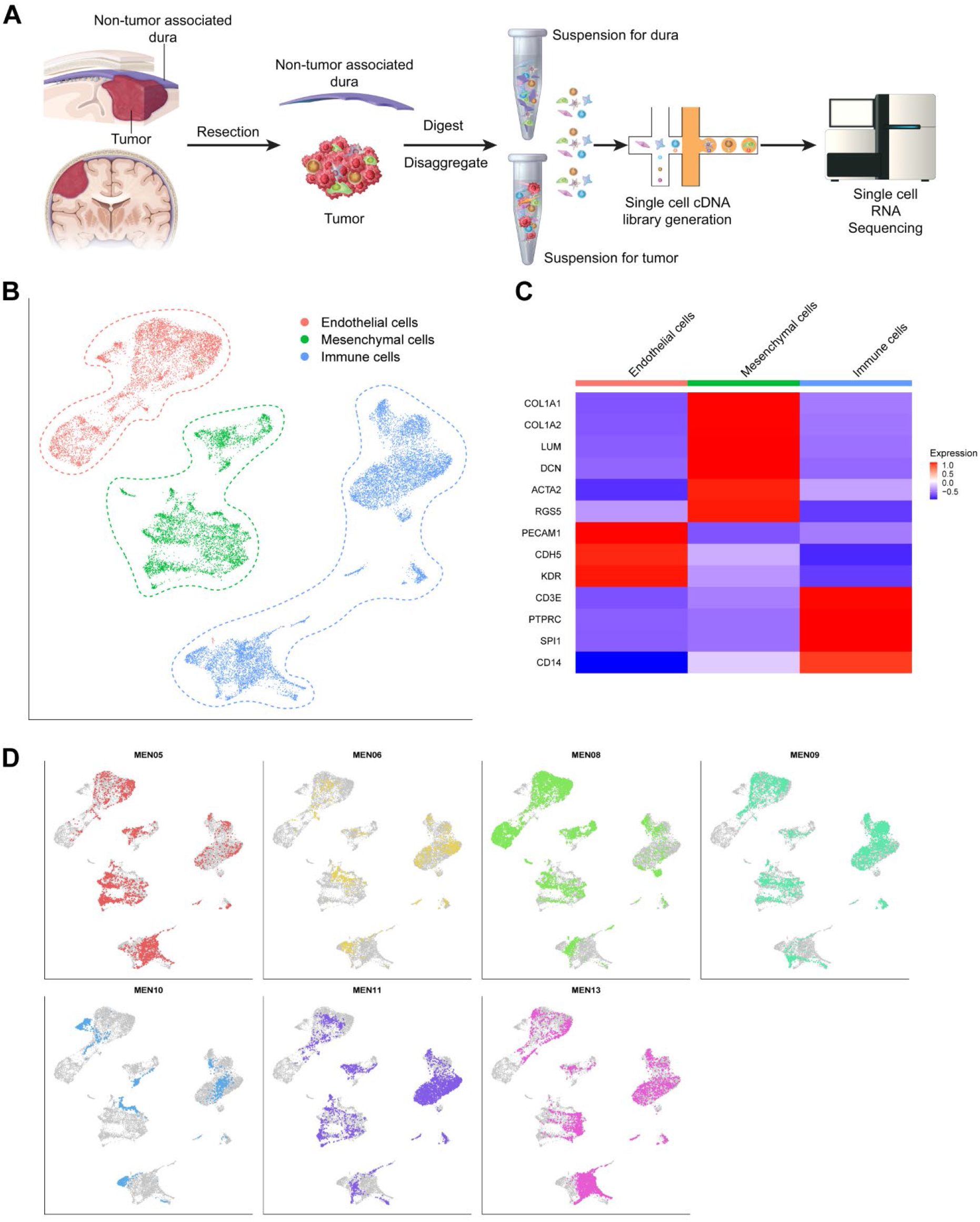
Single cell preparation and sequencing shows diverse cell landscape of human dura. **(A)** Illustration of both non-tumor associated dura and tumor resection and single cell library preparation. **(B)** UMAP visualization of single-cell RNA-seq data identified by cell type. **(C)** Representative gene expression of select cell type gene markers. **(D)** UMAP visualization of single-cell RNA-seq data highlighting cells originating from each individual patient sample.

### Immune cell composition of non-tumor-associated dura

Following a general analysis of the human dura samples, we next focused on resolving the immune cell (*PTPRC+*, which encodes CD45) landscape. To this end, we performed graph-based clustering and UMAP visualization on 10,423 immune cells (Fig. 2A). Clusters were characterized using a combination of previously reported marker gene sets and differentially expressed genes (Fig. 2B, Table S2, Data S1)(*16*–*25*). This revealed an appreciable population of lymphoid cells, including T cells, NK cells, B cells, and plasma cells, as well as myeloid cells, including monocytes, macrophages, DCs, and mast cells. UMAP visualization of important marker genes for lymphoid and myeloid cells are shown in indicated cell populations (Fig. 2C and 2D, respectively). T cells represented the majority of lymphoid cells observed (5,458 cells) and included naive/central memory (Tcm) cells (1,415 cells; which express *SELL, CCR7, LEF1, TCF7, KLF2*); CD4+ effector memory (Tem) cells (1575 cells; *SELL*-, *CCR7*-, *IL7R*); CD8+ Tem cells (1,281 cells; *SELL-, CCR7-, IL7R, CD8A, CD8B, GZMK*^*hi*^, *CXCR3*^*hi*^); resident memory (Trm) cells (251 cells; *CD69, NR4A2, IL7R*); and cytotoxic CD8+ T cells (936 cells; *PRF1, NKG7, ZNF683, GZMB, CD8A, CD8B*). Other lymphoid cell types identified included natural killer (NK) cells (571 cells; *PRF1, NKG7, GZMB, KLRD1, KLRF1, CD3-*), B cells (309 cells; *CD79A, MS4A1*, MHC class II+); and plasma cells (35 cells; *IGHG3, IGHA1, DERL1, FKBP11*). Meanwhile, the monocyte/macrophage/DC population (3,889 cells) was identified by monocyte-related (*CD14, VCAN, S100A8, S100A9*, MHC class II+), macrophage-related (*CD14, C5AR1, CD68, RNASE1, C1QC, CD163, FCAR, GPNMB*), and DC-related (*CD14-, ITGAX, THBD, and IL3RA*) markers. As the cell identities of these clusters were not clearly identifiable by gene marker sets at this resolution, we initially defined this population as being composed of monocytes, macrophages, and DCs. Finally, we identified mast cells (161 cells; *GATA2, KIT, HPGDS*) as an additional myeloid-based cell type within this population. These data demonstrate that, similar to murine dura, human dura is made up of a diverse population of immune cells.

**Figure 2:**
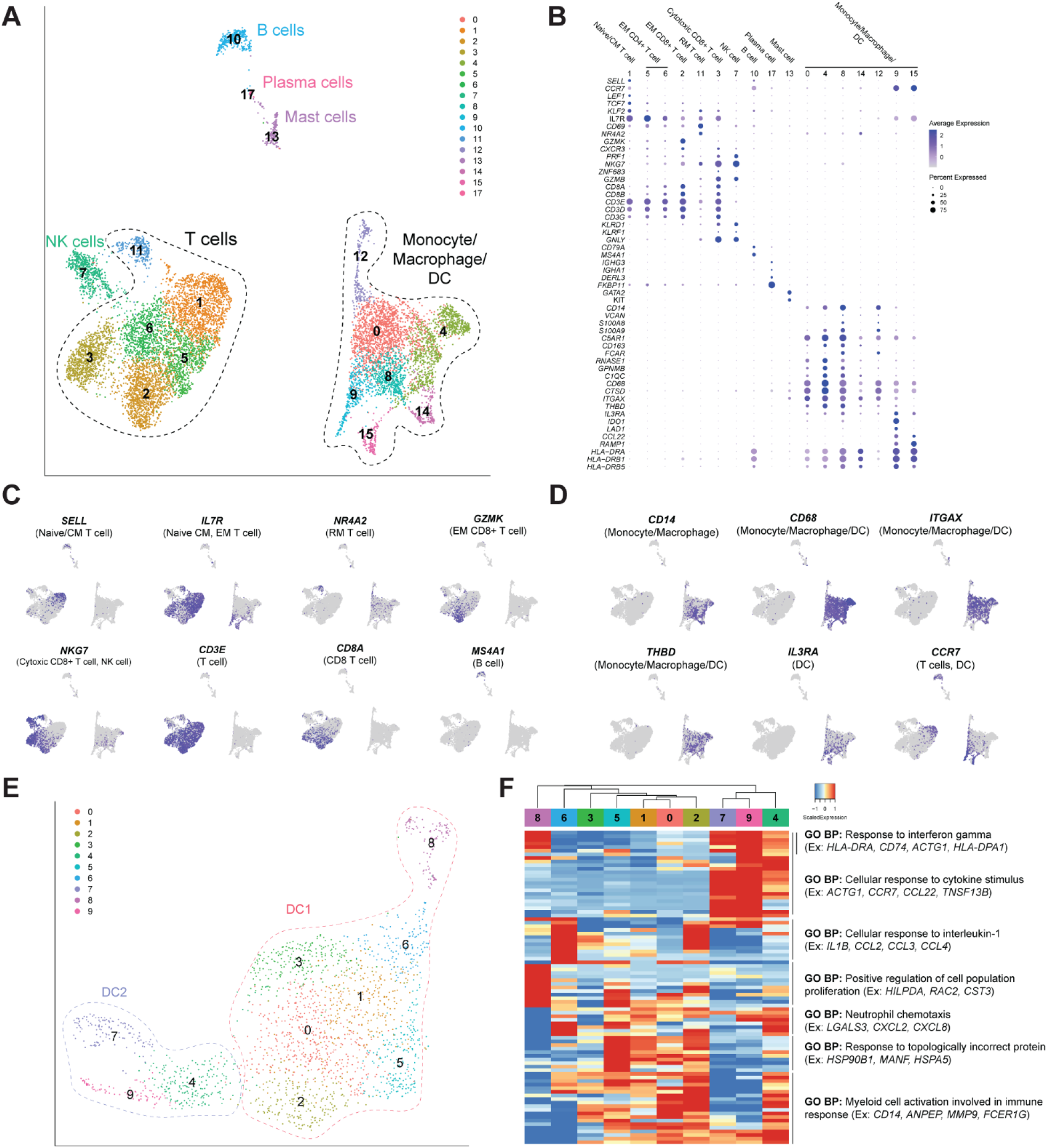
Immune cell composition of human dura consists of functionally diverse lymphoid and myeloid cell types. **(A)** UMAP visualization of immune cells identified by cell type (C1: Naive/CM T cells; C5, C6: EM CD4+ T cells; C2: EM CD8+ T cells; C3: Cytotoxic CD8+ T cells). **(B)** Representative gene expression of select cell type gene markers. **(C**,**D)** UMAP visualization of select lymphoid and myeloid marker gene expression. **(E)** UMAP visualization of dura DCs (C0, C1, C3, C4, C6, C7: DC1; C2, C5, C8: DC2). **(F)** Expression heatmap of top 10 genes of top 10 principal components with hierarchical clustering and associated functional enrichment analysis of gene clusters.

To better resolve the monocytes, macrophages, and DCs, we isolated all myeloid cells (excluding mast cells) for further analysis, including dimensionality reduction, clustering, and cell type annotation. We performed graph-based clustering and UMAP visualization on 3,889 cells (Fig. S1A), which revealed one monocyte/macrophage and two DC populations (DC1 and DC2) identified based upon marker gene sets (Table S2, Fig. S1B). Monocytes and macrophages were grouped together as these cells expressed varying levels of both marker gene sets. The low expression levels of microglial markers, such as *AIF1, C1QA*, and *GPR34*(*26*), and the expression of markers commonly associated with monocyte-derived macrophages suggested these cells were blood-derived rather than tissue resident. Interestingly, one cluster of cells (C6) was positive for monocyte and macrophage markers but lacked expression of MHC class II genes (*HLA-DRA, HLA-DRB1*), which suggested that they were myeloid-derived suppressor cells(*25*). Collectively, these data show myeloid cells occupy a considerable fraction of processed immune cells harbored by the dura.

We further characterized the DC population (DC1: *CD14-, ITGAX+ CD1c-, THBD*^int^; DC2: *CD14-ITGAX-IL3RA+ CCR7+*) using unsupervised clustering and UMAP visualization on 2,031 cells (Fig. 2E). To investigate variation within these cell populations, we applied hierarchical clustering and gene enrichment analysis using ToppGene(*27*) to the top 10 genes of the top 10 principal components (PCs) (Fig. 2F, Data S2). Our analysis indicated both the cell type present within the tissue as well as the biological pathways which associated with their respective gene expression profiles. DC1 clusters were characterized by the following pathways: “cellular response to interleukin-1,” “positive regulation of cell population proliferation,” and “response to topologically incorrect protein.” Meanwhile, DC2 clusters were characterized by the following pathway: “cellular response to cytokine stimulus.” Shared pathways included “response to interferon gamma,” “neutrophil chemotaxis,” and “myeloid cell activation involved in immune response.” Interestingly, Chen *et al*.(*20*) and Pombo Antunes *et al*.(*23*) observed DC clusters similar to DC2 that likewise differentially expressed genes such as *CCR7* and *LAMP3* and suggested them to be migratory DCs (migDCs) involved in immune cell recruitment. These data suggest that DCs harbored by the dura may be playing a role in establishing the dura immune microenvironment.

Collectively, our analysis demonstrated the presence of both lymphoid and myeloid cell subsets infiltrating the dura and underscored the dynamic nature of the dura immune microenvironment.

### Endothelial and mesenchymal cells comprise a significant proportion of cells in non-tumor-associated human dura

Having analyzed immune cells harbored by the dura, we next investigated non-immune (*PTPRC-*) cells by applying dimensionality reduction, clustering, and cell type annotation. Our analysis revealed three main cell types identified by previously reported marker gene sets and differentially expressed genes (Fig. 3A)(*28*–*31*): endothelial cells (6,157 cells; *PECAM1, CDH5, KDR, SELE, VWF*), fibroblasts (4,132 cells; *LUM, DCN, COL1A1, COL1A2, COL3A1*), and mural cells (1,368 cells; *ACTA2, MYH11, CNN1, RGS5, PDGFRB, NOTCH3, MCAM, CSPG4*) (Fig. 3B, Fig. 3C, Data S1). C18 (132 cells), an unidentified cell cluster, contained differentially expressed genes (DEGs) such as *TNNT3* and *DAT* (Data S1) not associated with any known common cell types. We also observed doublets (248 cells), defined by presence of immune related genes, though an increase in the number of genes was not detected. The presence of these cell types is consistent with our understanding of the gross structure of dura: a moderately vascularized tissue which harbors a collagen matrix scaffold which underpins the structure.

**Figure 3:**
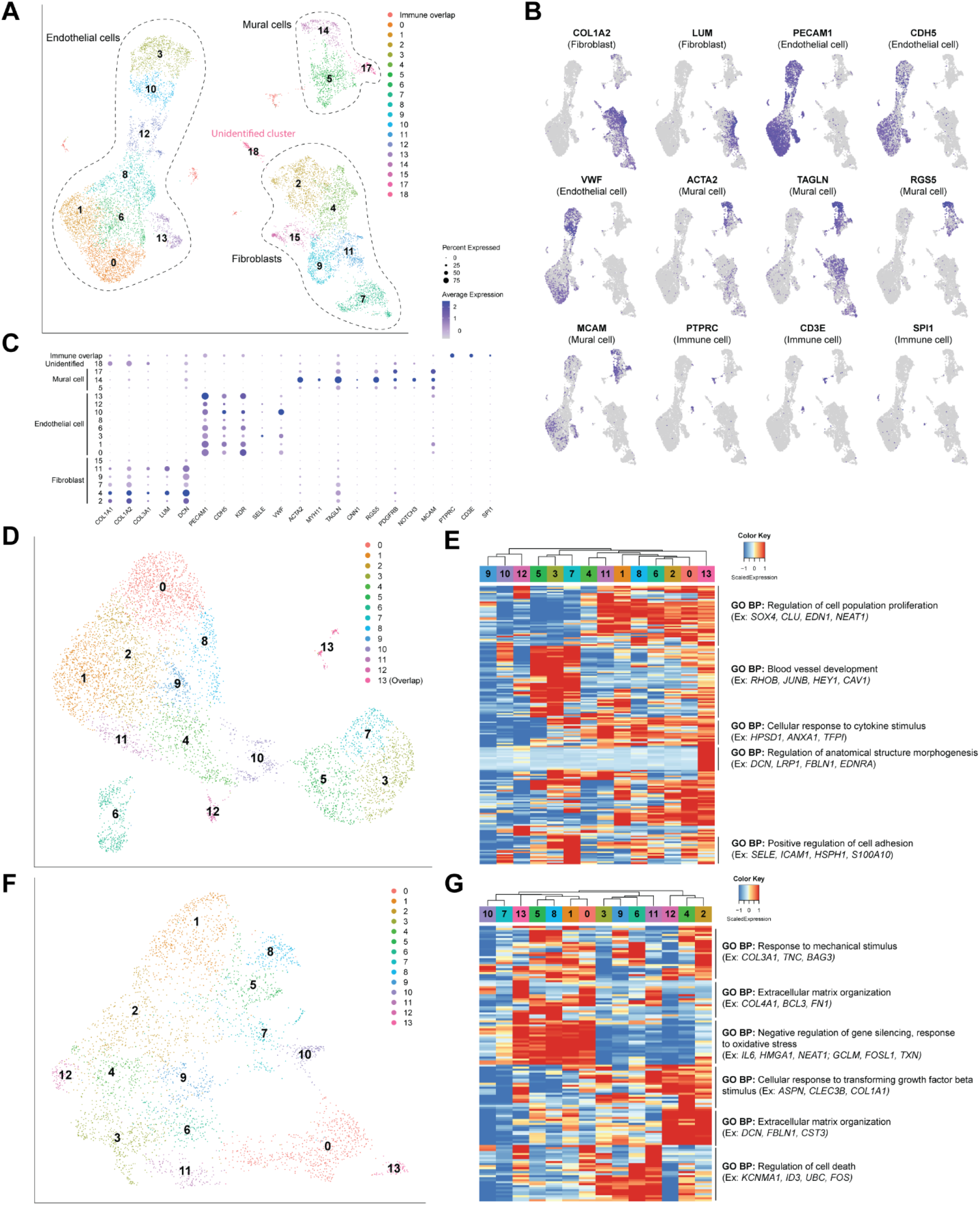
Functionally diverse non-immune cells comprise a significant proportion of human dura. **(A)** UMAP visualization of non-immune cells identified by cell type. **(B)** UMAP visualization of select endothelial, mural, and fibroblast markers. **(C)** Dotplot of representative gene expression of select cell type gene markers. **(D)** UMAP visualization of dura endothelial cells. **(E)** Expression heatmap of top 15 genes of top 15 principal components of dura endothelial cells with hierarchical clustering and associated functional enrichment analysis of gene clusters. **(F)** UMAP visualization of dura fibroblasts. **(G)** Expression heatmap of top 15 genes of top 10 principal components of dura fibroblasts with hierarchical clustering and associated functional enrichment analysis of gene clusters.

To better characterize the endothelial cell population, we selected and analyzed these cells using graph-based clustering and UMAP visualization (Fig. 3D). We hierarchically clustered the top 15 genes of the top 15 principal components to identify biological programs (Fig. 3E, Data S2). Endothelial cells fell into two main populations, with one distinguished by its expression of genes related to “regulation of cell population proliferation” and “cellular response to cytokine stimulus,” whereas the other was distinguished by “blood vessel development.” Both populations shared a biological pathway of “positive regulation of cell adhesion.” Notably, one cluster (C13) consisted of doublets due to expression of genes commonly associated with fibroblasts, such as *DCN*. From this analysis, we observed the majority of endothelial cells to be enriched for genes related to stimulus response and cell proliferation or blood vessel development. These data demonstrate evidence of a dynamic endothelial cell landscape in the dura layer composed of different endothelial cell populations with potentially unique functions.

We performed similar analysis on the fibroblast cell population by applying graph-based clustering and UMAP visualization (Fig. 3F). We analyzed the top 15 genes from the top 10 principal components (Fig. 3G, Data S2). In contrast to endothelial cells, fibroblasts exhibited considerably more heterogeneity in gene expression profiles and their associated biological pathways. As expected, the major pathway enriched by fibroblasts was “extracellular matrix organization,” though the genes which contributed to this biological pathway differed between two different populations of fibroblasts. Specialization of function was observed with one subset of clusters enriched for genes related to “negative regulation of gene silencing” and “response to oxidative stress,” while another was enriched for genes related to “regulation of cell death.” Finally, most clusters were enriched for genes related to “response to mechanical stimulus” and “cellular response to transforming growth factor beta.” “Response to oxidative stress,” “regulation of cell death,” and “response to mechanical stimulus” may, however, indicate cell stress due to the dissociation process rather than reflect the cell state within dura tissue. Similar to endothelial cells, fibroblasts in the dura are associated with many different biological functions though expression of said functions is more heterogeneously expressed among the fibroblast clusters.

These fibroblasts also harbored gene expression patterns similar to those of embryonic murine meninges fibroblasts recently described by DeSisto *et al*.(*32*). Specifically, we observed high expression of markers associated with murine dura fibroblasts defined by their experiments, such as *FXYD5, MGP, OGN*, and *TAGLN*, though not all dura fibroblast markers defined by their investigation, such as *TGFBI* and *CRABP2*, were highly expressed (Fig. S2). Arachnoid fibroblast markers, *ALDH1A2, WNT6*, and *CRABP2*, and pial fibroblast markers, *LAMA1* and *LAMA2*, were likewise expressed at low levels. Interestingly, *S100A6*, a pial fibroblast marker, was also highly expressed in these fibroblasts, similar to the unique M3-4 cluster of dura fibroblasts DeSisto *et al*. observed. Moreover, similar to a previous single-cell analysis by *Saunders et al*. of adult murine fibroblasts(*33*), we do not observe the expression of *DKK2* as seen in embryonic murine meninges fibroblasts by DeSisto *et al*. Thus, the similar gene expression patterns observed among embryonic and adult murine dura fibroblasts as well as our observations in adult human dura fibroblasts suggest cross-species conservation of fibroblast populations harbored by the dura.

### Imaging mass cytometry of human dura

Following characterization of human dura and meningioma at a single cell resolution, we performed imaging mass cytometry on available dura samples, DURA02 and DURA05, to visualize the spatial relationship among these cell types. Similar to our approach with scRNA-seq, we sought to first identify the immune, endothelial, and mesenchymal cell types using specific cell markers (Table S3). We identified vasculature by the presence of endothelial cells (CD31*+*, green) which was respectively surrounded by vascular smooth muscle cells (a-SMA+, red) in sample DURA02 (Fig. 4A). Moreover, we observed diffuse presence of collagen (cyan) throughout the tissue as expected. Next, we focused on immune cells, observing the presence of cells with overlapping expressions of CD14 and CD206, which may represent CD206^+^ macrophages (Fig. 4B). Meanwhile, we did not observe overlap between CD14 and Iba1 protein expression, a common microglial marker, suggesting a separate Iba1+ cell population is harbored by the dura (Fig. 4C). Within the dura, we also observed CD8+ T cells, represented by the overlap of CD3 and CD8a markers (Fig. 4D), though we did not observe CD4+ T cells, represented by the overlap of CD3 and CD4 markers (Fig. S3A). Surprisingly, we observed co-localization of CD8+ T cells and CD206+ cells outside CD31+ stained vasculature, rather than within, as denoted by the white arrow (Fig. 4E). In addition, we similarly observed co-localization of CD8+ T cells and Iba1+ cells (Fig. 4G). We also observed the presence of naive CD45RA+ T cells, denoted by overlap of CD45RA and CD3 (Fig. S3B), outside of CD31+ vasculature (Fig. 4F). Finally, we observed the presence of GZMB+ CD11b+ cells, often localized within CD31+ vasculature (Fig. 4H), indicating circulation of cytotoxic immune cells within non-tumor-associated dura. Imaging of DURA05 demonstrated similar results (Fig. 4I-N) with clear CD31+ vasculature surrounded by vascular smooth muscle cells (Fig. 4I) and the presence of CD8+ T cells (Fig. 4J) and prominent CD206+ and Iba1+ populations (Fig. 4L). Notable differences, however, included an abundant CD4+ T cell population (Fig. 4K), indicated by the white arrows, in addition to overlap between CD206 and Iba1 (Fig. 4L). We again observed co-localization between T cells, both CD4+ and CD8+ T cells, and CD206+ cells outside of CD31+ vasculature (Fig. 4M). Finally, we observed GZMB+/CD11b+ cells mostly within CD31+ vasculature though invasion beyond CD31+ vasculature was also noted (Fig. 4N). As co-localization of T cells and macrophages (represented by either CD206 or Iba1 expression) may suggest potential antigen presentation, we investigated whether genes associated with such pathways may be overrepresented in the single cell data. Specifically, applying CellChat(*34*), which infers and analyzes intercellular signaling pathways, to the immune cell population from Fig. 2A, we identified several signaling pathways that were significantly represented by the single cell data (Data S3). In particular, the monocyte/macrophage/DC population was the most prominent and significant source of MHC-I related ligands targeting the various T cell populations, with the exception of resident memory T cells (left side of Fig. 4O). Some autocrine signaling was observed with EM CD4+ T cells, EM CD8+ T cells, and cytotoxic CD8+ T cells. Furthermore, as expected, no significant relationships were observed with non-T cells as the target (right side of Fig. 4O). While these data raise the possibility that interaction between APCs and T cells may occur within the dura tissue itself, further investigation will be required to fully understand the functional roles of these immune cells within the dura as our current conclusions are limited due to the low number of samples and sites of imaging.

**Figure 4:**
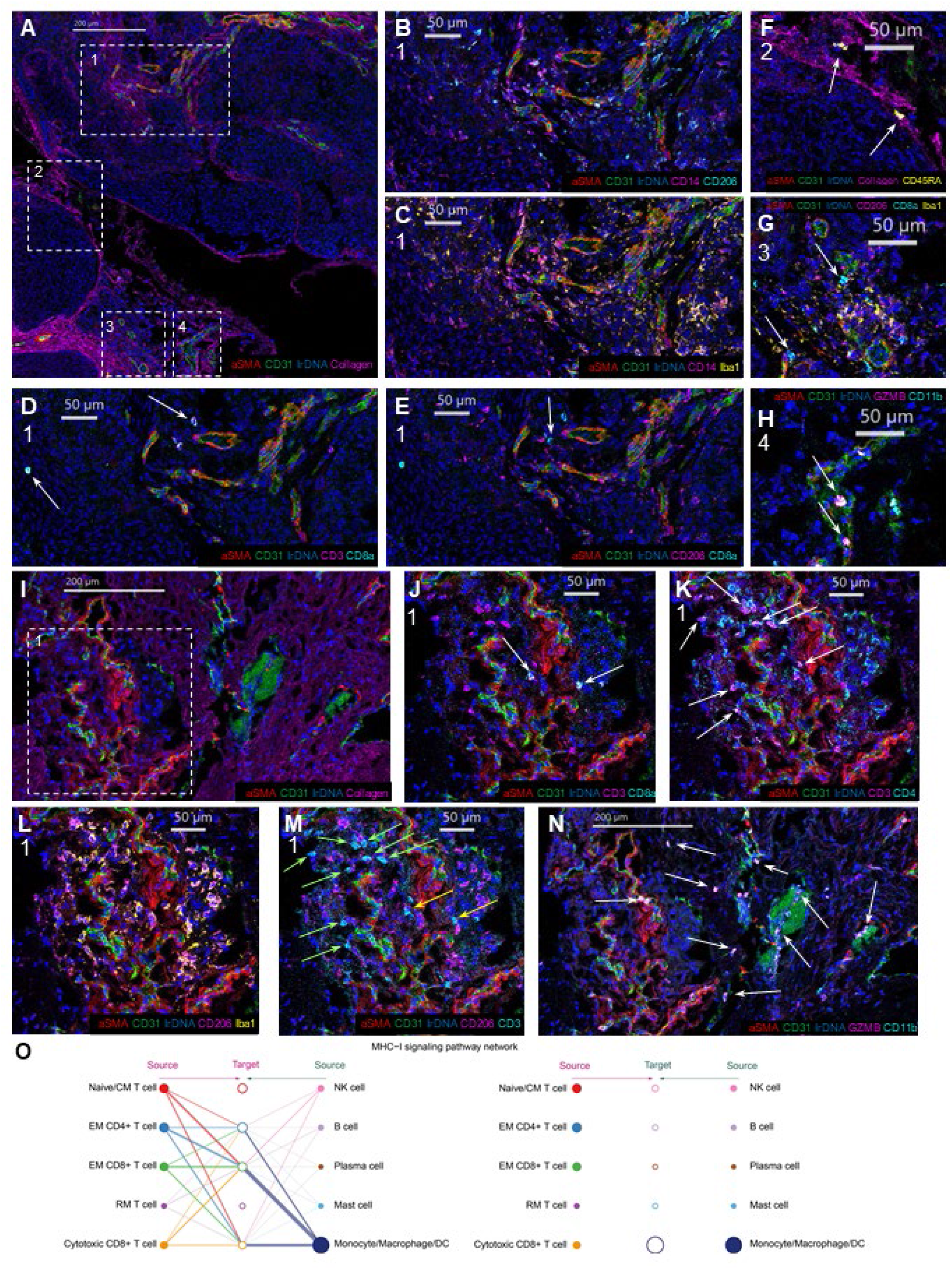
Imaging mass cytometry of a human dura sample reveals intricate spatial relationships among immune, endothelial, and mesenchymal cell types. **(A)** Imaging mass cytometry of human dura sample DURA02 labeled with markers specified and specific regions of interest (ROIs) highlighted by dashed white boxes. **(B-H)** Relative position of each image is denoted by marked number below panel letter. White arrows label cells of interest. **(I)** Imaging mass cytometry of human dura sample DURA05 with markers specified and the specific region of interest (ROI) highlighted by a dashed white box. **(J-N)** Relative position of each image is denoted by marked number below panel letter, if applicable. White arrows label cells of interest, green arrows represent CD4+ T cells, and yellow markers present CD8+ T cells. **(O)** Hierarchical plot showing the inferred network for MHC-I signaling for immune cells from Fig. 2A. The left and right portions of the plot show autocrine and paracrine signaling to T cells and remaining immune cells, respectively. Solid circles represent the source of MHC class I ligands and open circles represent the target of said MHC class I ligands. Circle sizes are proportional to the number of cells and width of connecting lines indicate the communication probability of said interaction.

### Distinct gene expression profiles demarcate immune cells infiltrating meningiomas from those in non-tumor-associated dura

In a subset of patients in our cohort, we were able to collect matched meningiomas together with non-tumor-associated dura adjacent to the tumor (Table S1). We analyzed the immune cells of four paired meningioma and non-tumor-associated dura samples composed of 12,581 cells and used the same markers described above to identify cell types. Within each cell type, we observed clear differences in cell state between cells isolated from each location (Fig. 5A, Data S1, Table S2, Fig. S4). Notably, we observed T cells, NK cells, monocytes/macrophages/DCs, and mast cells cluster separately based on tissue origin whereas B cells from both dura and tumor clustered together. Though dura T cells consist of naive T cells, Tem cells, and cytotoxic CD8+ T cells, only Trm cells were observed in the tumor samples.

**Figure 5:**
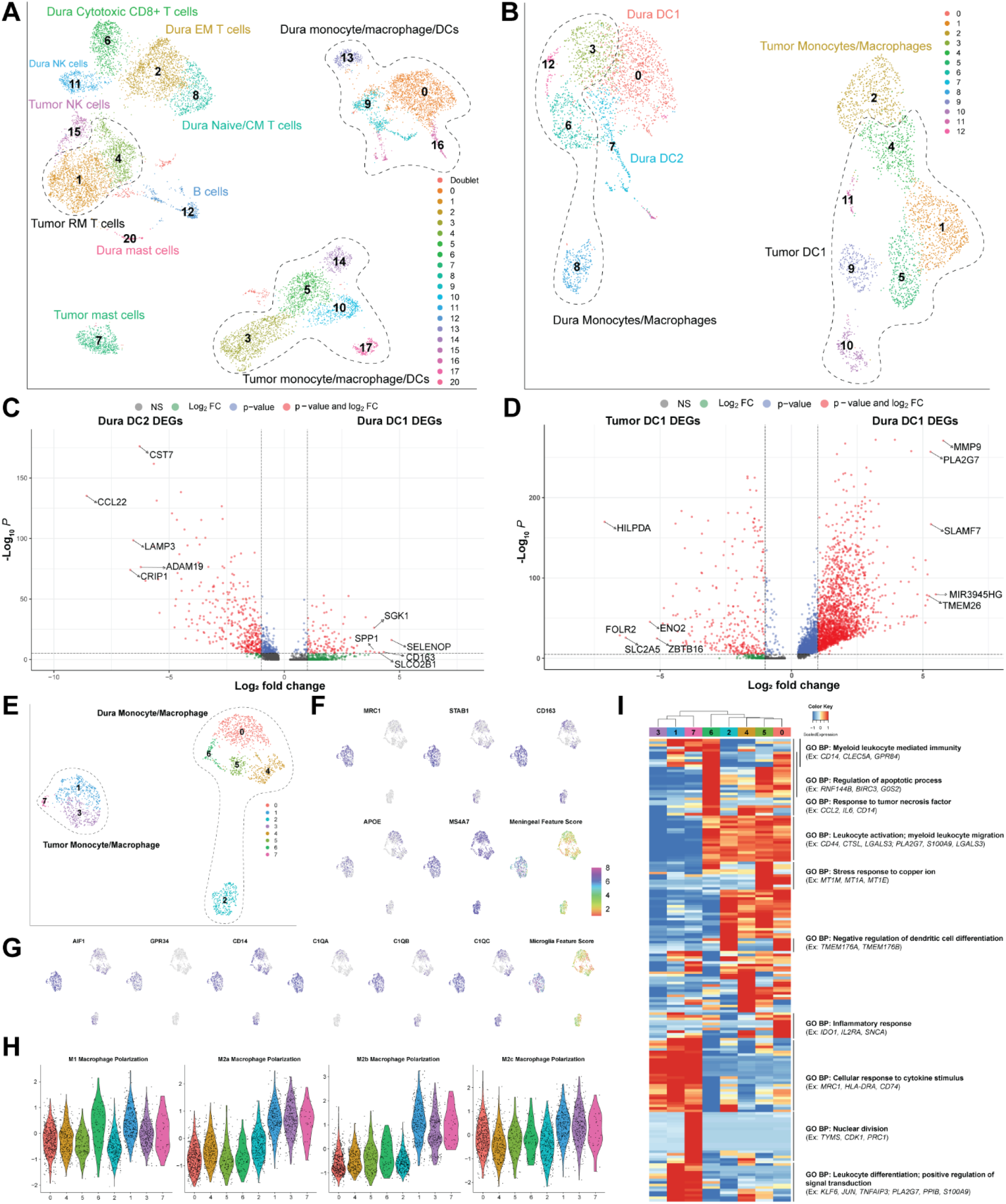
Normal human dura and meningioma samples show distinctively different immune cell populations. **(A)** UMAP visualization of dura and tumor immune cells identified by cell type. **(B)** UMAP visualization of dura and tumor monocyte/macrophage/DCs identified by cell type. **(C)** Volcano plot of genes differentially expressed by dura DC1 (positive log2 fold change) and DC2 (negative log2 fold change). **(D)** Volcano plot of genes differentially expressed by dura (positive log2 fold change) and tumor (negative log2 fold change) DC1. Log2 fold change and p-value thresholds are >|2| and 10e-6, respectively. **(E)** UMAP visualization of dura and tumor monocyte/macrophages identified by cell type. **(F)** UMAP visualization of select meningeal gene markers and aggregated score. **(G)** UMAP visualization of select microglial gene markers and aggregated score. **(H)** Violin plots of M1, M2a, M2b, and M2c macrophage polarization state scores of each cluster from Fig 4E. **(I)** Expression heatmap of the top 15 genes of the top 15 principal components with hierarchical clustering and associated functional enrichment analysis of gene clusters.

Comparing top DEGs which differentiate dura-originating from tumor-originating T cells, dura T cells DEGs were related to T cell migration and function, such as *CXCR3*(*35*) and *ITGAL*(*36*), as well as cell motility genes *SUSD3* and *FGD3*(*37, 38*) (Data S4). In contrast, tumor T cells differentially expressed genes coding for heat shock proteins, such as *HSPA6, HSPA1A*, and *HSPA1B* in addition to genes related to T cell development and function, such as *NR4A1*(*39, 40*) and *NR4A2*(*39*). Tumor NK cells expressed similar DEGs to tumor T cells and were enriched for genes associated with protein folding and cytokine expression, including genes coding for heat shock proteins *HSPA6, HSPA1B*, and *HSPA1A*, and *IFNG*, a common cytotoxic marker (Data S4). However, dura NK cells were enriched for genes that are associated with NK effector function, such as *SH2D1B*(*41*) and *KLRF1*(*42*). Interestingly though, *NLRC3*, a negative regulator of the innate immune response(*43*), was also overexpressed. Collectively, these data suggest that T cells and NK cells might have different functions in immune regulation depending on tissue of residence. However, as mentioned previously, though both types of tissues were processed similarly, presence of heat shock proteins may indicate differing responses to the dissociation process rather than reflecting differing cell states within respective tissues. Further investigation will be required to elucidate the clinical implications of these differences.

To further explore the differences among dura and tumor monocytes, macrophages, and DCs, we isolated and reanalyzed both dura and tumor myeloid clusters (excluding mast cells) (Fig. 5B). Marker gene sets were used to differentiate monocyte/macrophages, DC1, and DC2 cell clusters (Table S2, Fig. S5). Both dura and tumor DCs were separated and reanalyzed (Table S2, Fig. S6A, Fig. S6B). Pairwise comparisons of gene expression profiles were performed for dura DC1 and DC2 and dura and tumor DC1 clusters (Fig. 5C and 5D). Notable top DEGs expressed in dura DC1, compared to dura DC2, include *CD163*, an anti-inflammatory marker expressed by IL-10-producing DCs(*44*), *SLCO2B1*, a gene associated with DCregs(*45*), *SGK1*, and *SPP1*, which encodes for OPN and is implicated in DC maturation in addition to T cell polarization(*46*). Meanwhile, top DEGs expressed in dura DC2 include *CST7*, which has been shown to be upregulated in the transition of monocytes to moDCs(*47*), *CCL22*, which has been shown to induce DC cellular contact with Tregs in murine lymph nodes(*48*), *LAMP3*, a DC maturation marker, *ADAM19*, and *CRIP1* (Data S5). When compared to tumor DC1s, dura DC1s were enriched for genes such as *MMP9*, which has been shown to be pivotal for DC migration(*49*), as well as other genes *PLA2G7, SLAMF7, MIR3945HG*, and *TMEM26*. In contrast, tumor DC1s highly expressed *HILPDA, FOLR2, SLC2A5, ENO2*, and *ZBTB16*. Whether these DEGs have additional functional implications requires further investigation. These data demonstrate that DC1 and DC2 may play different functional roles within the dura, and that DC1 might be performing differing functions in the dura compared to the tumor.

We next selected dura and tumor monocyte/macrophage clusters and reanalyzed them (Fig. 5E). We used markers associated with microglia and border-associated macrophages (BAMs) in murine models (Table S4) to determine the potential origin(s) of these tumor monocyte/macrophages(*6, 26*) (Fig. 5F, 5G). Both microglial and BAM markers were enriched in tumor only clusters, and not in dura only clusters, which suggests that these macrophages were tissue resident, and originated from either the dura or brain parenchyma rather than blood. However, further functional validation will be needed to determine the origin of these macrophages. Surprisingly, we observed a discrepancy between scRNA-seq and IMC data as within non-tumor-associated dura tissue, CD206+ cells and Iba1+ cells co-localized with CD8+ and CD4+ T cells via IMC (Fig. 4E, 4G, 4M). Meanwhile, our scRNA-seq data showed low expression of *MRC1* (which encodes CD206) and *AIF1* (which encodes Iba1) in non-tumor-associated dura. Potential reasons for these discrepancies include a difference in mRNA and protein levels, lack of sequencing depth to detect gene expression, and alterations in cell state due to sample processing. We were also interested in characterizing the potential functionality of these macrophages by first assessing the macrophage polarization states in both dura and tumor clusters. Previously reported markers were aggregated to generate scores for M1, M2a, M2b, and M2c polarization(*50*) (Fig. 5H, Table S4). Overall, we observed similar M1 and M2c scores between dura and tumor clusters. However, tumor clusters demonstrated markedly elevated M2a and M2b scores, which are associated with an anti-inflammatory immune environment. We also analyzed the top 15 genes of the top 15 principal components to infer biological pathways and determine whether they were distinct based upon tissue source (Fig. 5I, Data S2). Overall, we observed a differentiation in gene expression profiles between dura monocyte/macrophages and tumor macrophages with varying associated biological pathways. Most prominently, all dura clusters were enriched for genes associated with “leukocyte activation” and “myeloid leukocyte activation.” Specific dura clusters were enriched for genes associated with “regulation of apoptotic process,” “response to tumor necrosis factor,” “stress response to copper ion,” “negative regulation of dendritic cell differentiation,” and “inflammatory response.” Meanwhile, all tumor clusters were enriched for genes associated with “cellular response to cytokine stimulus,” with particular enrichment in MHC class II genes such as *HLA-DRA*. Particular tumor clusters were enriched for genes associated with “myeloid leukocyte mediated immunity,” “nuclear division,” “leukocyte differentiation,” and “positive regulation of signal transduction.” From this gene enrichment analysis, we observed a difference in gene expression profiles between dura and tumor clusters, in addition to a difference in associated biological pathways. General analysis of the monocyte/macrophage clusters in both dura and tumor demonstrated evidence of macrophages with different tissue origins and unique gene expression profiles and associated functional pathways. Collectively, these findings suggest that myeloid cells infiltrating the dura and the tumor play different functional roles depending on the tissue in which they reside.

### Single cell analysis demonstrates CNV heterogeneity in meningioma

In addition to analysis of the immune cells from paired dura and meningioma samples, we also performed copy number analysis on the non-immune cells from each dura and meningioma pair to identify putative tumor cells using the R-based package CONICSmat(*51*) (Fig. 6A). By characterizing these cells with respect to known chromosomal abnormalities observed in meningioma(*52*), we identified a population of cells within MEN09 harboring several chromosomal abnormalities, which we inferred to be tumor cells (Fig. 6B, Fig. 6C). Specifically, the chromosomes with the most prominent copy number variations were 1p, 1q, 6p, 6q, 9q, and 19q. Weak support for copy number variations on chromosomes 16q and 22q was also indicated by CONICSmat. As previously reported, deletion (del) of chromosome 1p and 6q and amplification (amp) of 1q and 9q are associated with meningiomas(*52*). Notably, we observed amplification of the 6p and 19q chromosomal arms even though 6p deletion is generally observed and loss of heterozygosity has been reported with 19q. We did not identify chromosomal abnormalities in other paired dura and tumor samples, however, reflecting the fact that our dissociation conditions may have compromised the tumor cells in those samples.

**Figure 6:**
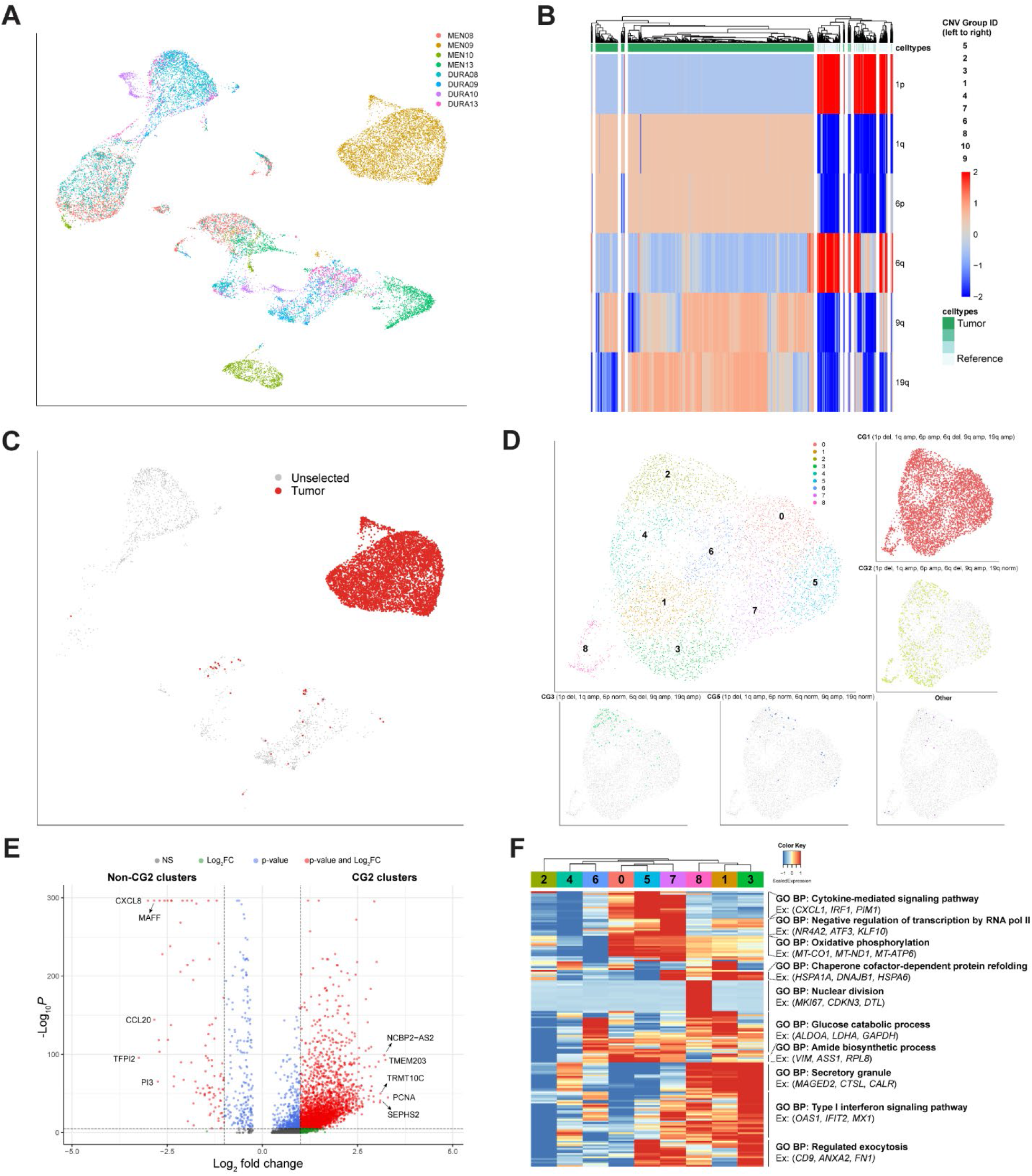
Analysis of meningioma cells reveals sub-clonal tumor populations with varying chromosomal abnormalities. **(A)** UMAP visualization of four paired dura and meningioma samples identified by sample origin. **(B)** Heatmap of DURA09 and MEN09 cells clustered by chromosome expression levels. **(C)** UMAP visualization of paired DURA09 and MEN09 samples with cell clusters that contain chromosomal abnormalities highlighted: CNV Group 1 (CG1), CG2, CG3, CG5, and Other. **(D)** UMAP visualization of meningioma cells with unsupervised clustering and highlighted by respective CNV group identity. **(E)** Volcano plot of genes differentially expressed by clusters containing CG2 (1, 2, 3, 4, 6, and 8) (positive log2 fold change) and non-CG2 clusters (negative log2 fold change). Log2 fold change and p-value thresholds are >|2| and 10e-6, respectively. **(F)** Expression heatmap of top 15 genes of top 15 principal components with hierarchical clustering and associated functional enrichment analysis of gene clusters.

Following our CNV analysis, we isolated, reanalyzed, and investigated these meningioma cells to better understand the CNV heterogeneity at the single cell level. Unsupervised clustering and UMAP analysis were performed on 6,287 cells in addition to visualization of specific groups of cells based upon degree of under- and overexpression of genes located on mentioned chromosomal arms (Fig. 6D). The major clonal population of cells belonged to CNV Group 1 (CG1) which contained 1p del, 1q amp, 6p amp, 6q del, 9q amp, and 19q amp relative to normal cells. In addition, we observed CG2 (1p del, 1q amp, 6p amp, 6q del, 9q amp, and 19q normal expression), CG3 (1p del, 1q amp, 6p norm, 6q del, 9q amp, 19q amp), and CG5 (1p del, 1q amp, 6p norm, 6q norm, 9q amp, and 19q norm) which may represent separate sub-populations. The remaining cells were categorized together and labeled as “Other”. Interested in further characterizing CG2 and CG3, we performed DEG analysis comparing clusters from Fig. 6D containing a significant number of CG2 and CG3 associated cells against clusters that did not, respectively (Fig. 6E, Fig. S7, Data S5). Genes enriched in clusters containing CG2 (C1, C2, C3, C4, C6, and C8) included *NCBP2-AS2, TMEM203, TRMT10C, PCNA*, and *SEPHS2. NCBP2-AS2* has previously been shown to be a regulator of angiogenesis following hypoxia in cancer-associated fibroblasts(*53*) and *SEPHS2* has been shown to be correlated with triple negative breast cancer aggressiveness and malignant tumor grade(*54*). However, *PCNA* is typically associated with vital cell processes such as DNA replication and repair of DNA damage(*55*). Meanwhile, genes enriched in clusters not containing CG2 (C0, C5, and C7) included *CXCL8, MAFF, CCL20, TFPI2*, and *PI3* (Fig. 6E, Data S5). *CXCL8* and *CCL20* have both been associated with the development of several cancer types(*56, 57*) while *TFPI2* and *PI3* downregulations are associated with tumor progression(*58, 59*). Genes enriched in clusters containing CG3 (cluster 2) included *FOXC1, JUND, NSMF, XIST, OBSCN* while genes enriched in clusters not containing CG3 included *ALDOA, IGF2, TM4SF1, IGFBP6*, and *PI3* (Fig. S7, Data S5). From these data, we identified significant differences in gene expression profiles of these various sub-populations.

Finally, we analyzed the top 15 genes of each of the top 15 PCs to identify expression programs within the clusters identified by unsupervised clustering as visualized in Fig. 6D (Fig. 6F, Data S2). Overall, several clusters showed an enrichment in genes related to metabolic pathways such as “oxidative phosphorylation,” “glucose catabolic process,” and “amide biosynthetic process,” thus indicating a metabolically active subset of meningioma cells. A subset of clusters was enriched for genes related to “cytokine-mediated signaling pathway,” “negative regulation of transcription by RNA pol II,” “oxidative phosphorylation,” “chaperone cofactor-dependent protein refolding,” “glucose catabolic process,” “amide biosynthetic process,” “secretory granule,” “Type I interferon signaling pathway,” and regulated exocytosis.” Notably, C2, which CG3 was primarily enriched in, expressed low levels of these top PC genes. C8 was distinguished from the other clusters by a significant enrichment in genes related to “nuclear division”. Thus, meningioma cells were differentiated by distinct gene expression and associated functional profiles though these profiles were not necessarily associated with specific CGs.

### TCR analysis of human dura and meningioma samples

To understand T cell clonotypic diversity within both matched dura and meningioma samples, we performed single cell sequencing on V(D)J region enriched libraries from four dura samples and two matched meningioma samples (Table S1). We first analyzed the relative frequency of T cell receptors (TCRs) by segregating the predominant clonotype (clone 1) from the rest, which were grouped based upon absolute count (i.e., clones 2-5, clones 6-20, clones 21-100, and clones 101-1000) (Fig. 7A). We observed a greater expansion of the top 20 clonotypes in the dura samples relative to those in the meningioma samples.

**Figure 7:**
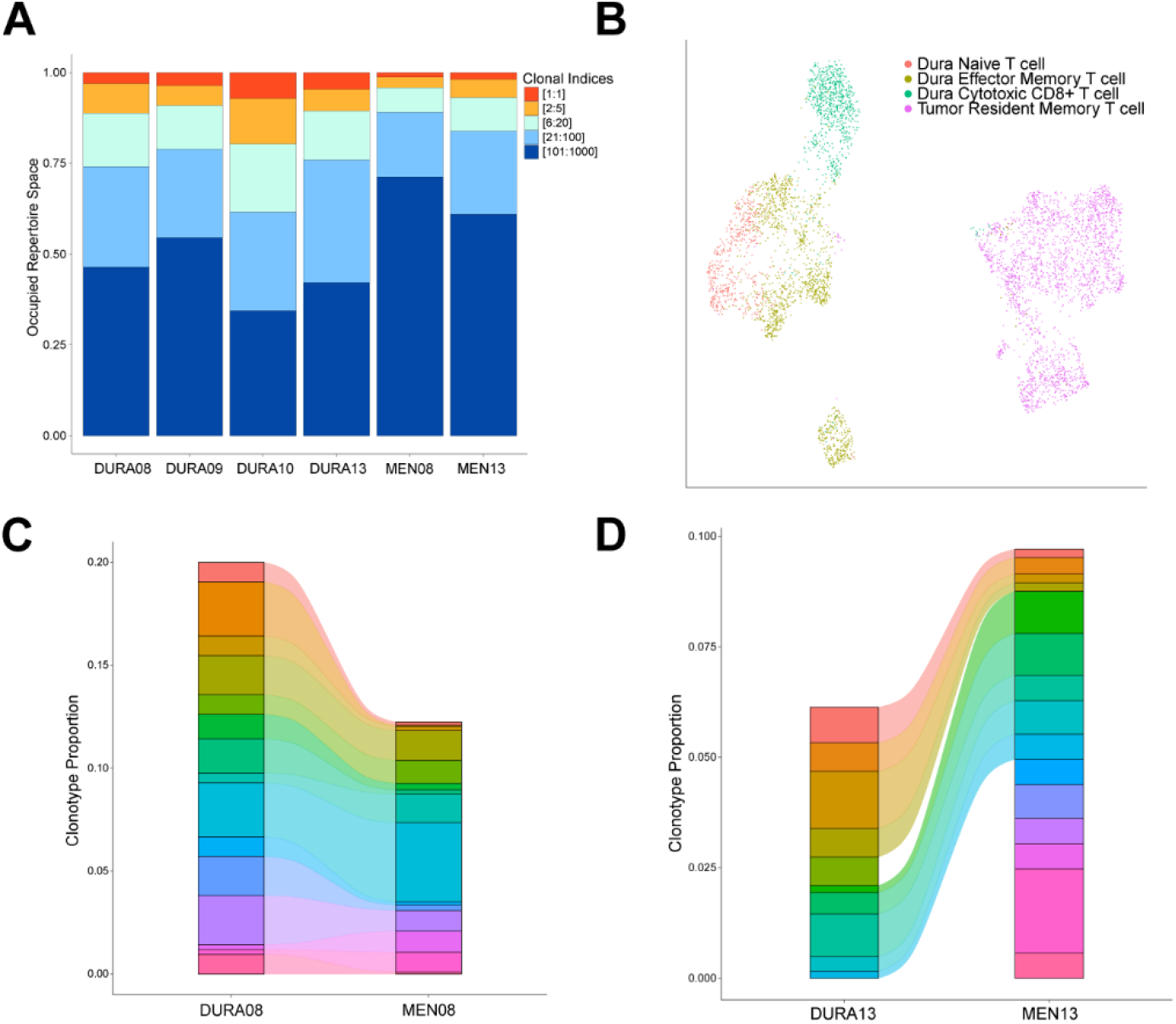
TCR frequency and expression overlap in paired dura and meningioma samples. **(A)** Clonal frequency of the dominant TCR, designated by absolute count, and groupings of TCRs ranked by absolute count. **(B)** UMAP visualization of dura and tumor T cells identified by cell type. **(C, D)** Alluvial plot demonstrating overlap of the top 15 and 16, respectively, TCRs in paired dura and meningioma samples ranked by relative frequency.

Following unsupervised clustering and UMAP analysis of SAMPLE08 and SAMPLE13 T cells alone from Fig. 5A, we observed a clear segregation of dura T cells from tumor T cells (Fig. 7B). Moreover, we generated alluvial plots of the top 15/16 TCRs, ranked by relative frequency with respect to each sample and represented by a distinct color, among the two paired dura and meningioma samples to determine the overlap of TCR presence (Fig. 7C and 7D, respectively). Strikingly, in the DURA08/MEN08 pair, all top 15 TCRs were identified at varying levels of expansion in both dura and tumor (Fig. 7C). Furthermore, DURA08 and MEN08 had a Morisita index, a measurement of the similarity between two data sets, of 0.484 when comparing the entire TCR repertoires of both samples, indicating a considerable amount of TCR overlap between the two samples (Fig. S8). In the DURA13/MEN13 pair, 9 of the top 16 most frequently expressed TCRs were present in both DURA13 and MEN13 samples (Fig. 7D). DURA13 and MEN13 have a Morisita index of 0.13, indicating a smaller, but non-zero, overlap of all TCRs compared to the MEN08 and DURA08 pair (Fig. S8). These data illustrate the T cell clonotypic diversity within the meninges and matched meningiomas and reveal that TCR clonotypes can be present within both meningiomas and nearby, non-tumor-associated dura sites. However, while CNV analysis supports that non-tumor-associated dura does not contain tumor cell clusters, our conclusions are limited due to the potential bias in tumor cell capture caused by sample preparation.

## Discussion

In this study, we present the first comprehensive single cell analysis of human non-tumor-associated dura and paired meningioma samples. The meninges has assumed growing importance in the study of CNS immunity and pathobiology as it has become clearer that it represents a dynamic microenvironment composed of unique cells with distinct immunologic, as well as non-immunologic, functions rather than a purely structural tissue barrier. To better understand this tissue site, we characterized both the immune and non-immune cell landscapes in non-tumor-associated dura in addition to analyzing the gene expression profiles of identified cell types using multiple platforms.

Building upon previous studies that characterized the immune microenvironment in murine dura(*6, 10, 60*), we observed a diverse collection of immune cells associated with both human non-tumor-associated dura and meningioma. The dura harbors several distinct T cell types ranging from T cells with a naive gene expression profile to those with cytotoxic profiles. The presence of these T cell subtypes may reflect ongoing immune surveillance by lymphocytes in non-tumor-associated dura, which has been demonstrated in murine models(*10, 60*). Meanwhile, the majority of T cells in meningioma samples were Trm cells, which play a pivotal role in protective immunity and have been identified in many human solid cancers and brain infection(*61*). Notably, the majority of Trm we observed were *CD8-*while current literature has focused on the role of *CD8*+ Trm in the immune response, especially against solid cancers. Similarly, we observed the presence of blood-derived monocyte/macrophage gene expression signatures in non-tumor-associated dura and macrophages enriched in border-associated macrophage (BAM) and microglial signatures, both of which are tissue-resident cell types, in tumor. However, from IMC of non-tumor-associated dura, we do detect the presence of cells expressing both BAM (CD206) and microglia (Iba1) related markers. This discrepancy may be due to lack of correlation between mRNA and protein expression, transcripts that are not detected by scRNA-seq (“dropouts”), and potential cell state changes due to sample processing. BAMs have been described to play immune roles, such as support and maintenance of barrier function and surveillance of antigens, within the meninges(*62*) and microglia have been implicated in brain parenchyma homeostasis(*63*). For these reasons, further investigation will be required to understand the function of these cells in non-tumor-associated dura compared to those present in the meningioma samples themselves. In addition, while the monocytes/macrophages from non-tumor-associated dura exhibited a pro-inflammatory gene expression profile, macrophages from tumor samples exhibited an anti-inflammatory expression profile, which has been previously observed in human meningioma samples via fluorescence immunohistochemistry(*64*). The significant difference in gene expression and associated functional profiles of immune cells based upon tissue origin warrants further investigation, as these differences may indicate targetable pathways to decrease tumor growth and the likelihood of recurrence.

Previous investigation in murine dura has established the role of the sinus vasculature in murine dura in allowing homeostatic T cell surveillance, in addition to demonstrating interaction between T-cells and APCs within the sinus and potential for T cells to infiltrate the meninges(*65*). Surprisingly, our imaging shows that such T cell and APC interactions may not be limited to the sinus region nor the vasculature, because co-localization of CD4+ T cells, CD8+ T cells, CD206+ cells, and Iba1+ cells was observed outside of CD31+ stained vasculature. In addition, we found within the single cell data that the monocyte/macrophage/DC population was a prominent source of MHC-I related ligands while the T cells were prominent sources of respective targets. As the antigens presented within the dura may differ from the CSF antigens presented by dura sinus-associated APCs(*65*), validation of such interactions and identification of presented antigens may present potential therapeutic targets.

By analyzing TCR clonotype diversity, we observed that the majority of highly expanded clonotypes were in cytotoxic CD8+ T cells collected from non-tumor-associated dura samples. Notably, tumor infiltrating T cells were less expanded and did not exhibit a robust cytotoxic CD8+ T cell phenotype, again illustrating a distinction between these environments though we are limited by the number of samples. In contrast, there are shared T cell clonotypes between paired non-tumor-associated dura and meningioma samples. Because meningiomas arise from the dura, these data suggest that tumor-specific T cells could enter meningiomas through the same blood vessels supplying the surrounding non-tumor-associated dura tissue(*66, 67*) following a priming event either within the dura or elsewhere. Furthermore, the presence of immune cell infiltrate in non-tumor-associated dura tissue observed via IMC and shared clonotypes in paired dura and meningioma samples suggest that non-tumor-associated dura may not be representative of normal dura in a healthy patient. In fact, our data suggests that an immune response may be occurring within the dura tissue itself. However, further investigation will be required to understand the mechanism behind T cell response to meningiomas, though the existence of shared T cell clones underscore the involvement of the dura in this process.

Meanwhile, investigating non-immune cells in human non-tumor-associated dura, we observed the presence of endothelial cells, fibroblasts, and mural cells, each of which plays an important role in the maintenance of the dura layer. The dura mater contains an abundant anastomotic arterial network(*68*) and in our samples, we observe an abundant presence of endothelial cells that are enriched in genes related to “blood vessel development” and “regulation of cell population proliferation”. The presence of mural cells, which have been shown to be present and regulate vascular diameter and blood flow in the CNS(*69*), is also observed in our samples. However, while the distinction between pericyte and vascular smooth muscle cell can be made based upon immunostaining techniques(*70*), this distinction is difficult to make based upon specific gene markers. As a result, further studies will be required to determine specific vasculature location and correlated gene expression profiles. Finally, fibroblasts have been shown to play an important role in CNS development(*1*) and may even contribute to nociceptive signaling in murine models(*71*). In our human dura samples, we observe an abundant presence of fibroblasts that contain a diverse gene expression profile enriched in biological pathways mainly concerning ECM development and response to various stimuli. These results indicate a dynamic cellular population in the human meninges, even beyond the development phase, that requires further investigation to better characterize and understand its role in maintaining dura mater homeostasis beyond a purely structural function. We also show cross-species conservation of dural specific fibroblast markers by comparing our data set to that of Desisto *et al*.(*32*) and Saunders *et al*.(*33*).

Finally, from our analysis of meningioma cells, we observe CNV heterogeneity with one major clonal population and three potential sub-populations each with different CNVs. Though the two larger sub-populations show CNV-associated gene expression profiles, they are not distinguished by specific biological pathways. Given our low number of samples taken at one time point, further studies are needed to understand the development of meningiomas and the relationship among these sub-populations and main clonal populations. In addition, with more samples, we can better determine the relationship among CNV heterogeneity, gene expression, and functional properties of meningioma cells. Interestingly, such transcriptomic intratumoral heterogeneity, in addition to epigenetic heterogeneity, has been associated with high grade meningiomas at a bulk-tissue level with accompanying single cell investigation of human cerebral organoids(*72*). However, beyond this, investigation into primary meningioma tumor samples at a single cell resolution has been relatively unexplored.

Our study provides the first scRNA-seq characterization of both immune and non-immune cell types in human non-tumor-associated dura and matched primary meningioma tumor samples. Similar to collaborative efforts such as the Human Cell Atlas(*73*), these data represent a resource to further understand the cellular composition of human dura. However, we recognize that there are several limitations of our study. First, given the proximity of non-tumor-associated dura to meningioma tumor, in addition to shared TCR clonotypes with paired meningioma samples, this dura may not be representative of truly “normal” tissue and thus additional opportunities to study dura in distinct clinical contexts will be important and necessary. Furthermore, we recognize the limited number of samples in our study, potential bias due to sample preparation, and limited validation of cell type presence via IMC. Given the growing evidence implicating the importance of the dura in biological pathways, such as immunosurveillance of the CNS, and in particular brain tumors(*13, 74*), and its current relevance in practical medical settings, such as embolization of the middle meningeal artery for new or recurrent chronic subdural hematoma(*14*), we envision there to be significant translational and clinical implications of an improved understanding of the biology of the dura.

## Materials and Methods

### Experimental Design

The objective of this study was to characterize the cellular composition of human dura and meningioma at a single cell resolution. The study design involved performing 3’ and 5’ single cell RNA-sequencing, with V(D)J enrichment for select samples, in addition to imaging mass cytometry.

### Sample extraction and preparation

Dura and matched tumor samples were collected from the operating room on ice. Excess sample was blotted dry and frozen in Fisher Healthcare Tissue-Plus Optimal Cutting Temperature (O.C.T.) Compound (Fisher Scientific) on dry ice and stored at -80°C. To disaggregate the remaining samples, samples were placed in a sterile-filtered medium containing 10%FBS (Lonza), IMDM (Lonza), 2 mg/mL collagenase A (Roche), and 2 mg/mL collagenase D (Roche), and macerated into small pieces with a scalpel. Samples were incubated overnight at 37°C with 5% CO2, with gentle pipetting intermittently to encourage further disaggregation. The following morning, after the collagen matrix had completely dissolved and the cells had dissociated into a single-cell suspension, the cell suspension was passed through a 100 µm strainer, followed by RBC lysis with ACK buffer (Lonza), followed by passing through a 70 µm strainer and then a 40 µm strainer. If a significant portion of the cells were dead, samples were subjected to EasySep™ dead cell removal per manufacturer’s protocol (Stemcell Technologies). If the suspension had considerable debris, samples were subjected to debris removal solution according to manufacturer’s instructions (Miltenyi Biotec). Cells were resuspended in 10% FBS/IMDM in preparation for construction of single-cell libraries.

### Single-cell RNA sequencing

Cell suspensions were prepared according to the manufacturer’s protocol (10x genomics) for 3’ v3 single cell sequencing or 5’ single cell sequencing with TCR enrichment (Table S1). For both methods, gel beads in emulsion (GEMs) were generated from a mixture of cell suspension combined with the GEM beads subjected to emulsion production by the Chromium Controller. cDNA was prepared after the GEM generation and barcoding, followed by the GEM-RT reaction and bead cleanup steps. Purified cDNA was amplified for 10-14 cycles before being cleaned up using SPRIselect beads. Samples were then run on a tape station or Bioanalyzer to determine the cDNA concentration. TCR enrichments were done on the full-length cDNA 5’ gene expression libraries (GEX). Both GEX and Enriched TCR libraries were prepared as recommended by the 10x Genomics Chromium Single Cell V(D)J Reagent Kits (v1 Chemistry) user guide with appropriate modifications to the PCR cycles based on the calculated cDNA concentration. For sample preparation on the 10x Genomics platform for 3’v3 libraries, the Chromium Single Cell 3’ GEM, Library & Gel Bead Kit v3 (PN-1000075) with the Chromium Single Cell Chip B Kit (PN-1000154) were used. For 5’ libraries the Chromium Single Cell 5’ Library and Gel Bead Kit (PN-1000006), Chromium Single Cell A Chip Kit (PN-1000152), Chromium Single Cell V(D)J Enrichment Kit, Human, T cell (96 rxns) (PN-1000005), and Chromium Single Index Kit T (PN-1000213) were used. The concentration of each library was accurately determined through qPCR utilizing the KAPA library Quantification Kit according to the manufacturer’s protocol (KAPA Biosystems/Roche) to produce cluster counts appropriate for the Illumina NovaSeq6000 instrument. Normalized libraries were sequenced on a NovaSeq6000 S4 Flow Cell using the XP workflow and a 151×10×10×151 sequencing recipe according to manufacturer protocol for 5’ sequencing and a 28×8×98 sequencing recipe according to manufacturer protocol for 3’v3 sequencing. For both sequencing approaches, a median sequencing depth of 50,000 reads/cell was targeted for each Gene Expression Library and 5000 reads/cell for each V(D)J (Tcell) library generated from the 5’ sequencing library.

### Single-cell RNA-seq data processing of dura and meningioma samples

Raw sequencing data was processed with the CellRanger pipeline (10x Genomics, default settings, Version 3.0.1) mapped onto a human genome GRCh38-3.0.0. All seven dura samples were then processed using the Seurat R package(*75*) and cells that contained fewer than 500 features, more than 10% mitochondrial transcripts, and a nCount value greater than the 93rd percentile of each individual sample were removed. Cells containing greater than 6,000 nFeatures was removed for DURA08 sample. Samples were then batched according to sequencing technology (3’ or 5’ sequencing) and each batch was individually log normalized after which variable features were selected according to default settings. Both batches were then integrated using FindIntegrationAnchors and IntegrateData. Principal component analysis was then performed and the optimal number of principal components (PCs) was determined based upon results from the elbow plots, jackstraw resampling, and PC expression heatmaps (n=50). Dimensionality reduction and visualization were performed with the UMAP algorithm (Seurat implementation) and unsupervised graph-based clustering was performed at a resolution of 0.7. Cell cycle phase was assessed based upon expression of phase-specific genes following methodology provided by Seurat(*76*).

Similarly, four paired dura and meningioma samples were processed together using the Seurat R package. Similar filtering criteria were applied however samples were merged, rather than batched and integrated as the same sequencing technology was used, and normalized. Variable features were selected according to default settings and principal component analysis was performed. The optimal number of principal components was determined (n=50) and dimensionality reduction and visualization were performed with the UMAP algorithm. Unsupervised graph-based clustering was performed at a resolution of 0.7 and cell cycle phase was similarly assessed.

Differentially expressed genes of each cluster resolved by unsupervised graph-based clustering were determined using a Wilcoxon Rank Sum test-based function. These genes, along with commonly defined markers (Table S2), were used to identify cell identity.

Based upon cell type classification (i.e. immune, non-immune, monocyte/macrophage, etc.), clusters of cells were individually isolated after which each data set was rescaled and variable genes were again selected according to default settings. Principal component analysis was performed and an appropriate number of principal components selected (dura immune cells: n=30, dura myeloid cells: n=20, dura DCs: n=15, dura non-immune cells: n=30, dura endothelial cells: n=20, dura fibroblasts: n=20, dura and tumor immune cells: n=30, dura and tumor myeloid cells: n=15, dura and tumor DCs: n=15, dura and tumor macrophages: n=15, dura and tumor non-immune cells: n=30, and tumors cells: n=20). Dimensionality reduction and visualization were performed with the UMAP algorithm and unsupervised graph-based clustering was performed at the following resolutions (dura immune cells: 1.0, dura myeloid cells: 0.6, dura DCs: 0.9, dura non-immune cells: 0.9, dura endothelial cells: 0.9, dura fibroblasts: 0.7, dura and tumor immune cells: 0.7, dura and tumor myeloid cells: 0.8, dura and tumor DCs: 0.9, dura and tumor monocyte/macrophages: 0.6, dura and tumor non-immune cells: 0.8, and tumor cells: 0.7).

### Immunohistochemistry validation of antibodies and antibody conjugation

Purchased antibodies (Table S3) were initially tested via immunohistochemistry. Respective positive control tissues for each antibody were tested (Novus Biologicals). Slides were first baked in an oven at 56 °C overnight to melt the paraffin wax, then placed in xylene for 20 minutes, and rehydrated in the following metal-free solutions of ethanol for 5 minutes each: 100%, 100%, 95%, 95%, 80%, 80%, 70%, 70%. After rehydration, slides were placed in metal-free water for 5 minutes on an orbital shaker after which they were incubated in pH 9 IHC Antigen Retrieval Solution (Invitrogen) at 96°C for 30 min. Slides were cooled in the antigen retrieval solution for 10 minutes at room temperature, followed by a 10-minute wash in metal-free water and a 10-minute wash in metal-free PBS. The tissues on the slides were outlined with a hydrophobic barrier pen (Liquid Blocker) and a solution of 3% bovine serum albumin (BSA) in PBS was placed on the tissues within the hydrophobic barriers for 45 minutes at room temperature. The primary antibody solution was prepared at manufacturer-recommended dilutions in PBS with a final concentration of 0.5% BSA and subsequently added following removal of the 3% BSA solution. The primary antibody solution was incubated on the tissues overnight at 4 degrees Celsius. Slides were placed within a hydration chamber during this time. Following overnight incubation, the slides were then washed in 0.2% Triton-X 100 in PBS for 10 minutes twice and washed in PBS for 10 minutes twice. Secondary antibody solution was prepared similar to the primary antibody solution with secondary antibody diluted following manufacturer recommendations in PBS with a final concentration of 0.5% BSA. Slides were incubated with secondary antibody solution away from light for 45 minutes at room temperature and then washed in 0.2% Triton-X 100 in metal-free PBS for 10 minutes, twice and then in metal-free PBS for 10 minutes, twice. Slides were then treated with 3 uM DAPI solution for 2 minutes away from light and subsequently washed in metal-free PBS for 10 minutes. Tissues were then mounted with coverslips using Vectashield Plus Antifade Mounting Medium (Vector Laboratories) and sealed with clear nail polish (Revlon 771). Tissues were imaged on Zeiss LSM 880 with oil immersion.

Antibodies with successful positive staining were subsequently conjugated to lanthanide metals (Table S3) following the protocol associated with the Maxpar X8 Antibody Labeling Kit (Fluidigm).

### Imaging Mass Cytometry

Dura and tumor samples frozen in O.C.T. at -80°C were thawed, removed from O.C.T., and immediately fixed in 10% neutral buffered formalin for 22 hours. Samples were then transferred to 70% metal-free ethanol, embedded in paraffin wax, and sectioned at a thickness of 5 um onto standard slides. Slides were baked in an oven at 56°C overnight to melt the paraffin wax and then placed in xylene for 20 minutes and then rehydrated in the following metal-free solutions of ethanol for 5 minutes each: 100%, 100%, 95%, 95%, 80%, 80%, 70%, 70%. After rehydration, slides were placed in metal-free water for 5 minutes on an orbital shaker and then incubated in pH 9 IHC Antigen Retrieval Solution (Invitrogen) at 96°C for 30 min. Slides were cooled in the antigen retrieval solution for 10 minutes at room temperature and washed in metal-free water for 10 minutes and metal-free PBS for 10 minutes. The tissues on the slides were outlined with a hydrophobic barrier pen (Liquid Blocker) and a solution of 3% bovine serum albumin (BSA) in PBS was placed on the tissues within the hydrophobic barriers for 45 minutes at room temperature. The lanthanide-conjugated antibody solution, with respective dilutions as outlined in Table S3, was prepared in PBS with a final concentration of 0.5% BSA. This antibody solution was placed on the tissues within the hydrophobic barrier following removal of the 3% BSA solution. Slides were incubated with this antibody solution overnight at 4°C in a hydration chamber. Following overnight incubation, the slides were then washed in 0.2% Triton-X 100 in PBS for 8 minutes, twice. They were then washed in PBS for 8 minutes, twice. DNA-Intercalator (Fluidigm) solution was prepared in PBS at a dilution of 1:400 and incubated on the tissues within the hydrophobic barrier for 30 minutes at room temperature. Slides were washed in metal-free water for 5 minutes and then air-dried for 20 minutes. The prepared slides were imaged on the Hyperion System (Fluidigm). Imaging results were visualized through MCD Viewer (Fluidigm) and saved as 16-bit TIFF images. Individual channel intensities were manually selected and standardized throughout all images.

### Expression heatmaps & gene functional enrichment analysis

Expression heatmaps were generated by selecting the top selected number of genes in each of the top selected number of PCs. The expression of each gene was averaged within each cluster and scaled and the results were hierarchically clustered using heatmap2. Gene functional enrichment analysis was performed using ToppGene (https://toppgene.cchmc.org/enrichment.jsp)(*27*). Hierarchically clustered gene groups were selected and the top one or two gene ontology biological pathways were displayed. All gene groups are listed in Data S2.

### Macrophage polarization, meningeal macrophage, and microglial scores

Macrophage polarization, meningeal macrophage, and microglial scores were generated using *AddModuleScore* (Seurat implementation) and previously published gene lists(*6, 26, 50*).

### TCR analysis

Raw TCR sequencing data was processed with the Cellranger V(D)J pipeline (10x genomics, default settings, Version 2.0.0) mapped onto a human VDJ reference GRCh38-2.0.0. Clonotype analysis was performed using the scRepertoire R package(*77*). All data processing was performed as outlined [here](https://ncborcherding.github.io/vignettes/vignette.html). The following clonotype states were as defined: hyperexpanded (50 < X <= 150), large (20 < X <= 50), medium (5 < X <= 20), small (1 < X <= 5), and single (0 < X <=1).

### Copy number analysis

Copy number alterations were assessed using the CONICSmat package for R(*51*). Gene expression values were filtered and normalized as discussed [here](https://github.com/diazlab/CONICS/wiki/Tutorial---CONICSmat;---Dataset:-SmartSeq2-scRNA-seq-of-Oligodendroglioma). The following chromosomal positions were the final ones assessed: 1p, 1q, 6q, and 9q. The z-score posterior probabilities were clustered, with a cut-off score of z=2, and cell barcodes from the ten clusters were gathered and visualized on UMAP.

## Statistical analysis

Differential gene expression was calculated using the Wilcoxon rank-sum test (two-sided) as implemented in the Seurat R package. Bonferroni correction was used to adjust *p*-values based upon total number of features in the dataset. Enriched gene ontology biological pathways were assessed using ToppGene and the top one or two biological processes significant after Bonferroni correction were selected.

## Acknowledgments

Immunohistochemical slides of dura were imaged on a Zeiss LSM 880 Airyscan Confocal Microscope which was purchased with support from the Office of Research Infrastructure Programs (ORIP), a part of the NIH Office of the Director under grant OD021629, and imaging was performed in part through the use of Washington University Center for Cellular Imaging (WUCCI) supported by Washington University School of Medicine, The Children’s Discovery Institute of Washington University and St. Louis Children’s Hospital (CDI-CORE-2015-505 and CDI-CORE-2019-813) and the Foundation for Barnes-Jewish Hospital (3770 and 4642). This work was also supported, in part, by the Bursky Center for Human Immunology and Immunotherapy Programs at Washington University, Immunomonitoring Laboratory.

We thank the Genome Technology Access Center at the McDonnell Genome Institute at Washington University School of Medicine for help with genomics services. The Center is partially supported by NCI Cancer Center Support Grant #P30 CA91842 to the Siteman Cancer Center and by ICTS/CTSA Grant# UL1TR002345 from the National Center for Research Resources (NCRR), a component of the National Institutes of Health (NIH), and NIH Roadmap for Medical Research. This publication is solely the responsibility of the authors and does not necessarily represent the official view of NCRR or NIH.

AZW was supported by the Medical Scientist Training Program at Washington University in St. Louis and JABK was supported in part by the Medical Scientist Training Program at Washington University in St. Louis and in part by a F30 grant.

We finally would like to thank the Arthritis & Clinical Immunology Human Phenotyping Core at the Oklahoma Medical Research Foundation for assistance with Hyperion Imaging Mass Cytometry data acquisition.

## Author contributions

Conceptualization: JABK, AAP, GPD

Methodology: JABK, BP, PRP, DB, MCM

Investigation: AZW, JABK, BP, RD, JWO, ECL, MRC, RGD, GJZ, AHK

Formal analysis: AZW, SMK, JL, AAP

Software: AZW, SMK, JL, AAP

Resources: RD, JABK, BP, AAP, GPD

Supervision: AAP, GPD

Validation: AZW, DB, AAP, GPD

Visualization: AZW

Writing-original draft: AZW

Writing-review and editing: AZW, JABK, RD, BP, AAP, GPD

Funding acquisition: GPD

Data curation: AZW, AAP

Project administration: RD, AAP, GPD

## Competing interests

GPD is a member of the Scientific Advisory Board of Ziopharm Oncology, the clinical advisory board of ImmunoGenesis, and is a co-founder of Immunovalent.

AHK has received research grants from Monteris Medical for a mouse laser therapy study as well as from Stryker and Collagen Matrix for clinical outcomes studies about a dural substitute, which have no direct relation to this study.

ECL has listed the following competing interests:

**Acera**:

- Compensation: Service on an Advisory Board: ($0)
- Equity in non-publicly traded entity:

**Alcyone**:

- Compensation: Consulting (not otherwise specified):

**Caeli Vascular**

- Equity in non-publicly traded entity: ($1-$4,999)
- Licensing/Product Development Agreements or Royalties for inventions/IP:

**Cerovations**

- Licensing/Product Development Agreements or Royalties for inventions/IP:

**E15**

- Compensation: Consulting (not otherwise specified):

**Immunovalent**

- Equity in non-publicly traded entity:

**inner Cosmos**

- Equity in non-publicly traded entity:

**Intellectual Ventures**

- Compensation: Consulting (not otherwise specified):
- Licensing/Product Development Agreements or Royalties for inventions/IP:

**Kinexus**

- Equity in non-publicly traded entity:

**Monteris**

- Compensation: Consulting (not otherwise specified):

**Neuroloutions, Inc**.

- Compensation: Consulting (not otherwise specified):
- Compensation: Service on a Board of Directors:
- Equity in non-publicly traded entity:
- Licensing/Product Development Agreements or Royalties for inventions/IP:

**Osteovantage, Inc**.

- Compensation: Service on an Advisory Board:
- Equity in non-publicly traded entity:
- Licensing/Product Development Agreements or Royalties for inventions/IP:

**Pear Therapeutics, Inc**.

- Compensation: Service on an Advisory Board:
- Equity in non-publicly traded entity:

**red devil 4**

- Equity in non-publicly traded entity:

**Sante Ventures**

- Compensation: Consulting (not otherwise specified):

**Sora Imaging Solutions**

- Equity in non-publicly traded entity: ($0)
- Licensing/Product Development Agreements or Royalties for inventions/IP:

**SympEL Neuromodulation**

- Equity in non-publicly traded entity:

All other authors declare they have no competing interests.

## Data and material availability

Single cell RNA-sequencing (scRNA-seq) data generated during the current study are available on the open-access data sharing platform Zenodo (http://doi.org/10.5281/zenodo.5120927).

