## Supplementary Tables and Figures for "Single Cell Atlas of Human Dura Reveals Cellular Meningeal Landscape and Insights into Meningioma Immune Response"

**This PDF file includes:**

Figs. S1 to S8
Tables S1 to S4
Data S1 to S5

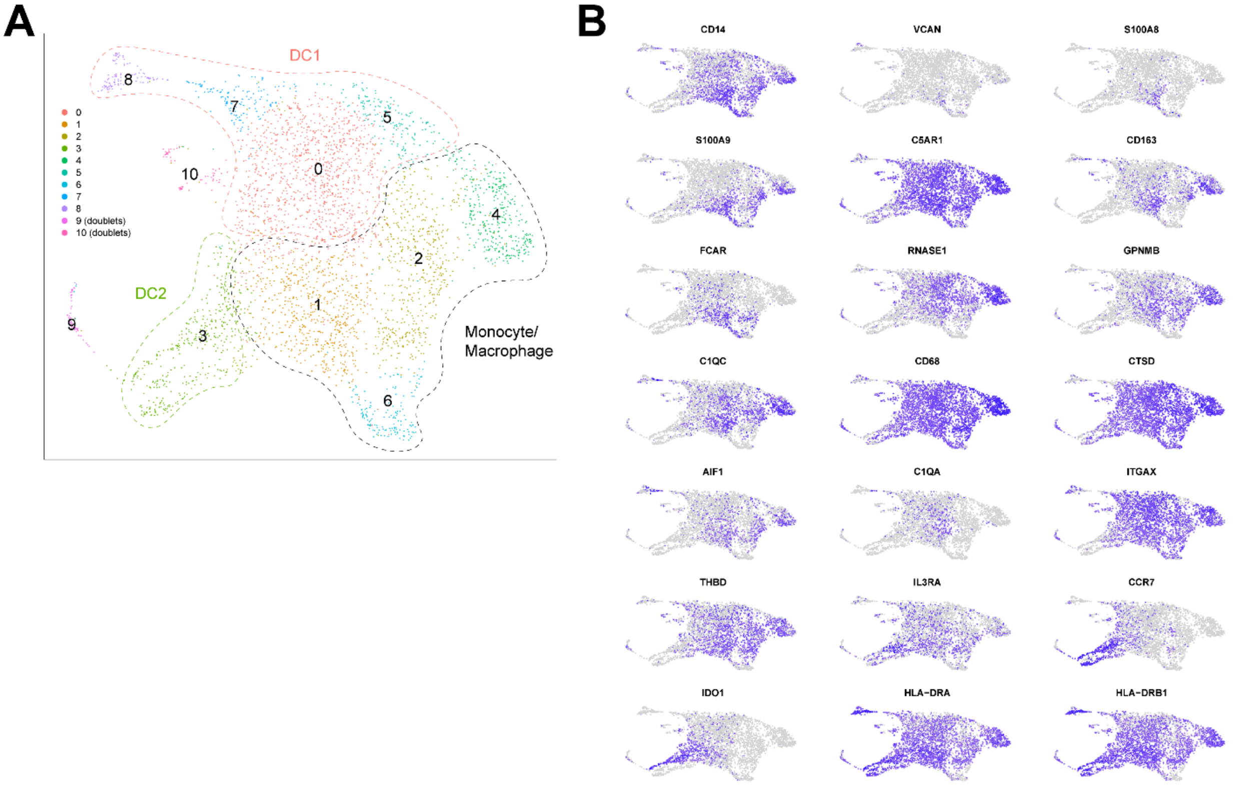


**Fig. S1. Analysis of dura myeloid cells excluding mast cells. (A)** UMAP visualization of dura myeloid cells excluding mast cells. Monocyte/macrophages include C1, C2, C4, and C6. DC1 includes C0, C5, C7, and C8. DC2 includes C3. **(B)** UMAP visualization of dura myeloid cells excluding mast cells for select monocyte, macrophage, and DC marker genes.

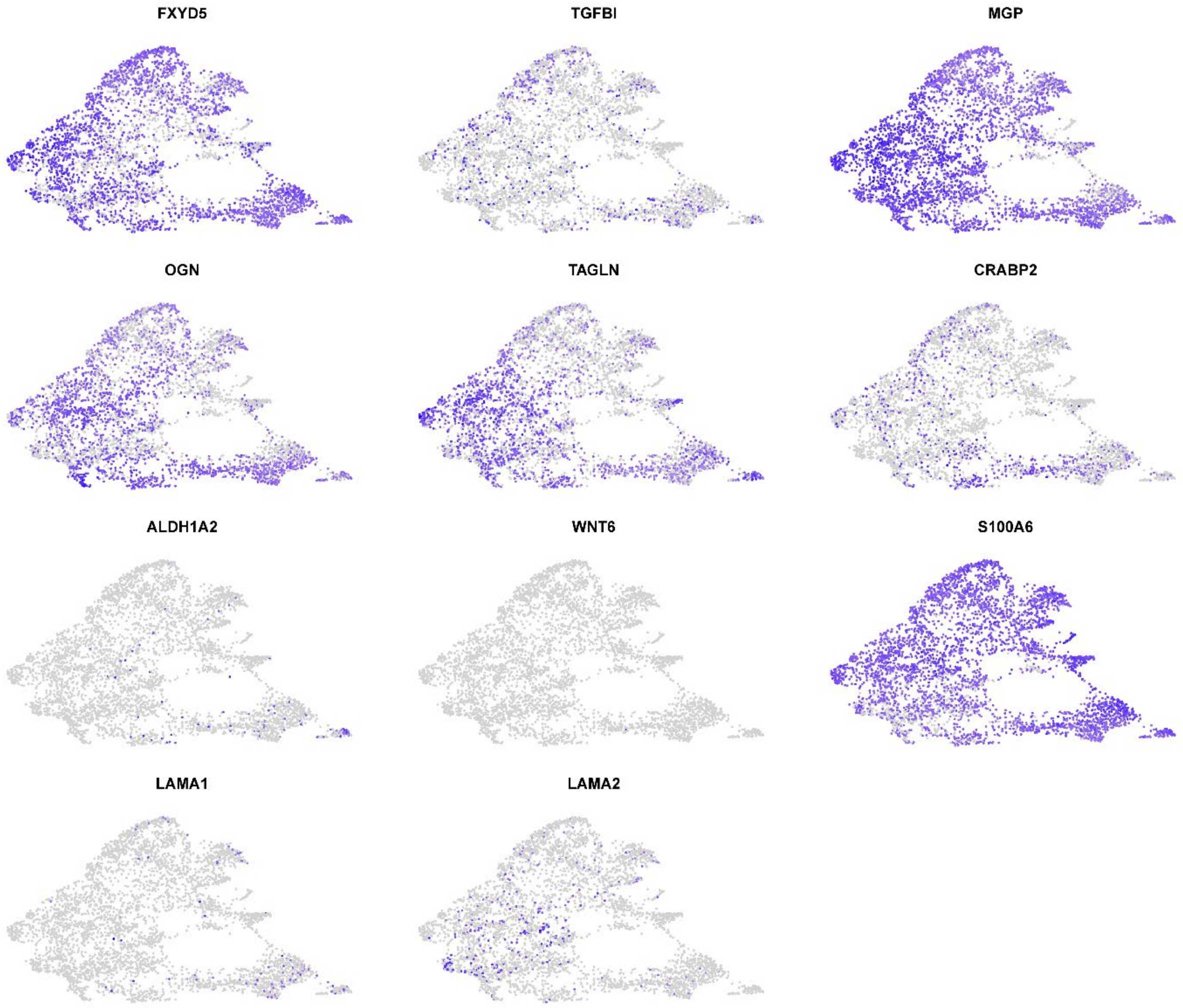


**Fig. S2. Meningeal fibroblast markers.** UMAP visualization of *CD45*- cells in non-tumor associated dura for specific meningeal fibroblast markers as discussed by DeSisto *et al.* (31) (dura fibroblasts markers: *FXYD5, TGFBI, MGP, OGN, TAGLN, CRABP2*; arachnoid fibroblast markers: *CRABP2, ALDH1A2, WNT6, TAGLN, OGN*; pia fibroblast markers: *S100A6, LAMA1/2, CRABP2-, ALDH1A2-*).

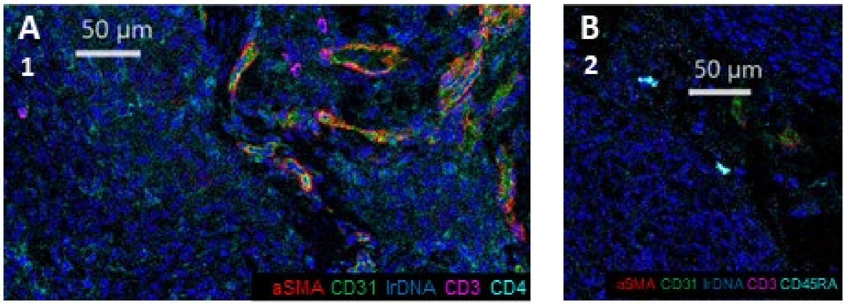


**Fig. S3. Imaging mass cytometry of DURA02. (A, B)** Imaging mass cytometry of human dura sample DURA02 labeled with markers specified. Relative position of each image is denoted by marked number below panel letter in reference to marked positions in Fig. 4A.

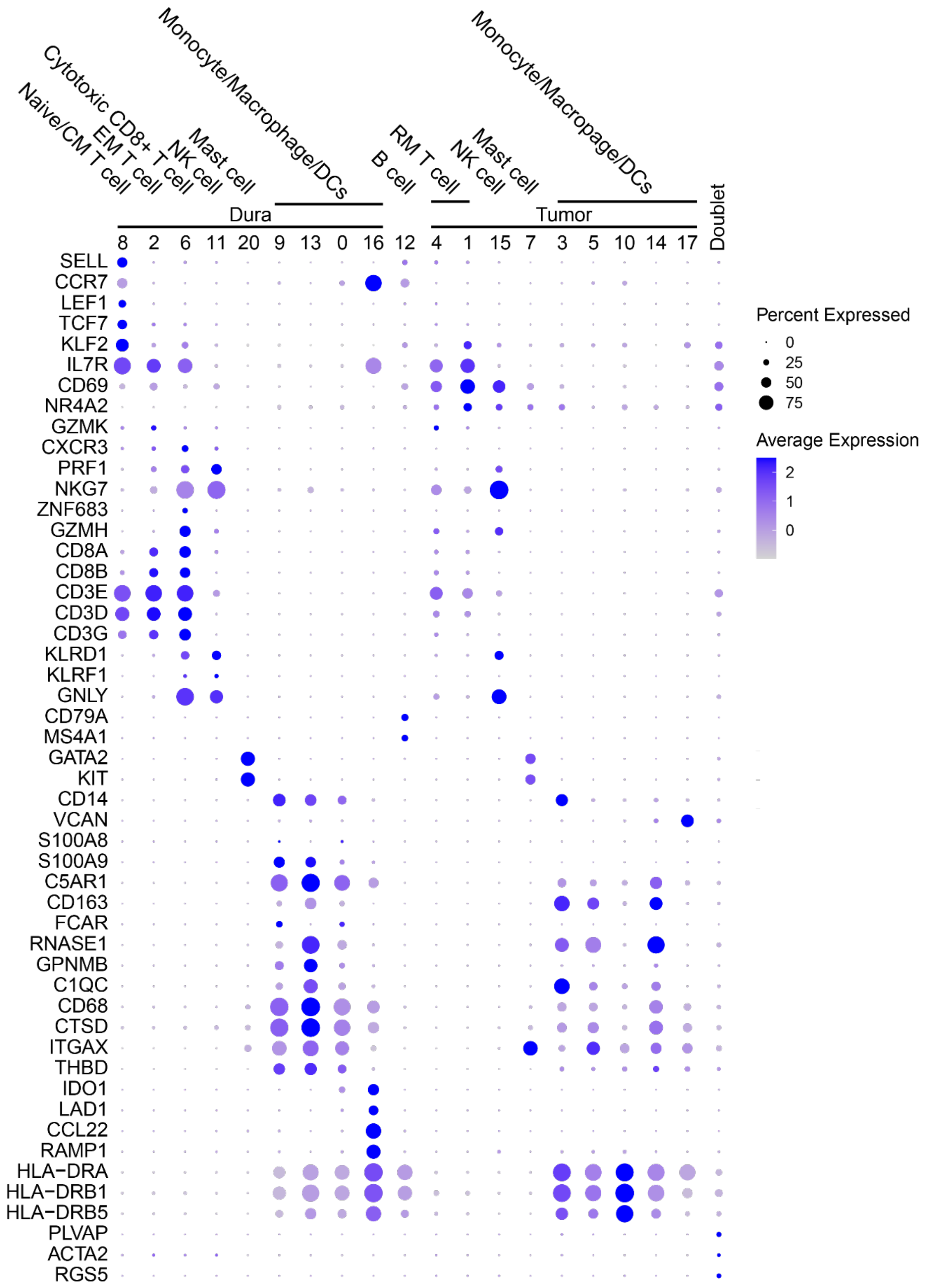


**Fig. S4. Gene markers for *CD45*+ cells in dura and tumor samples.** Dot plot visualization of gene markers used for cell identity identification of *CD45*+ cells in dura and tumor samples.

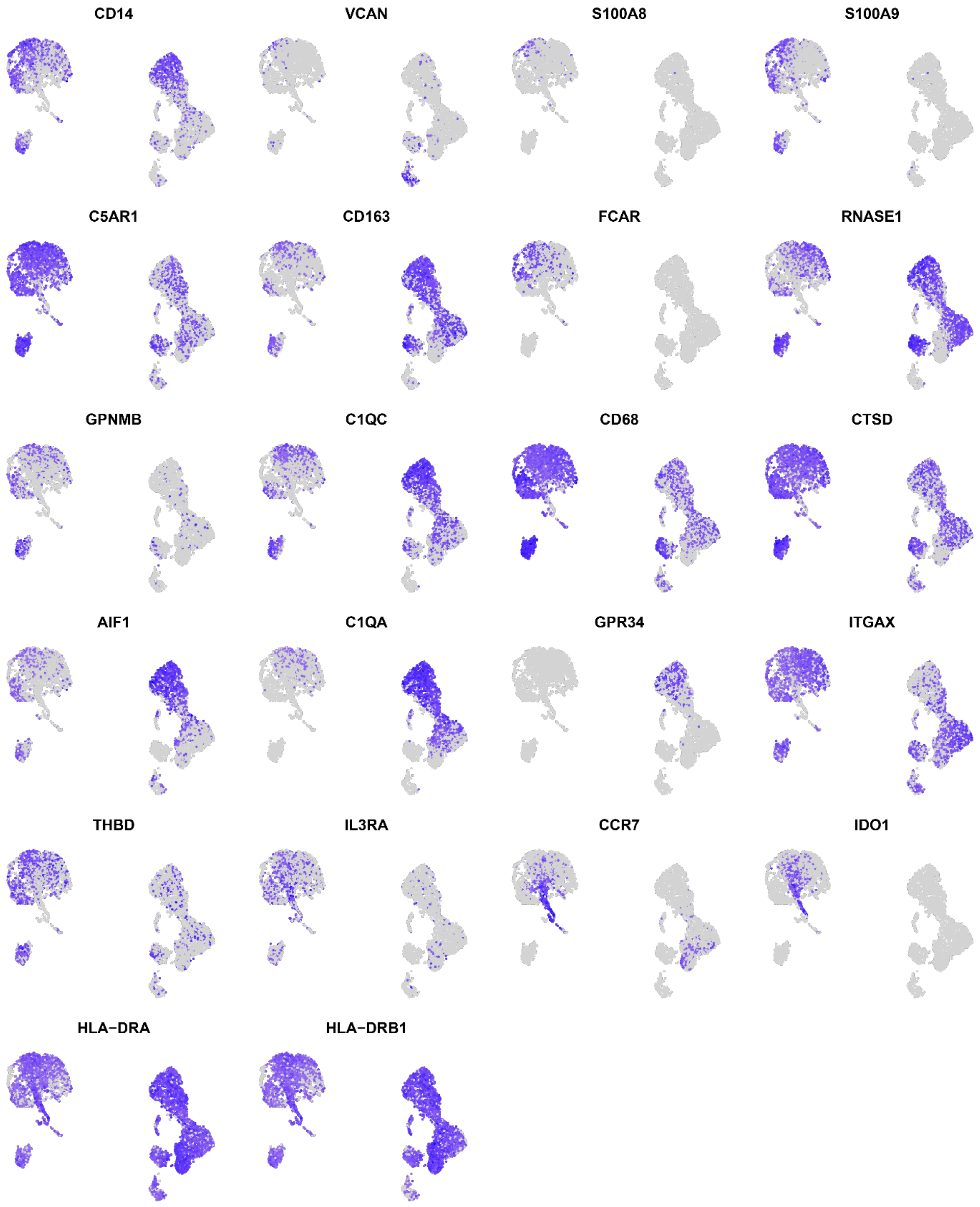


**Fig. S5. Marker genes of dura and tumor myeloid cells.** UMAP visualization of dura and tumor myeloid cells for select monocyte, macrophage, and DC marker genes.

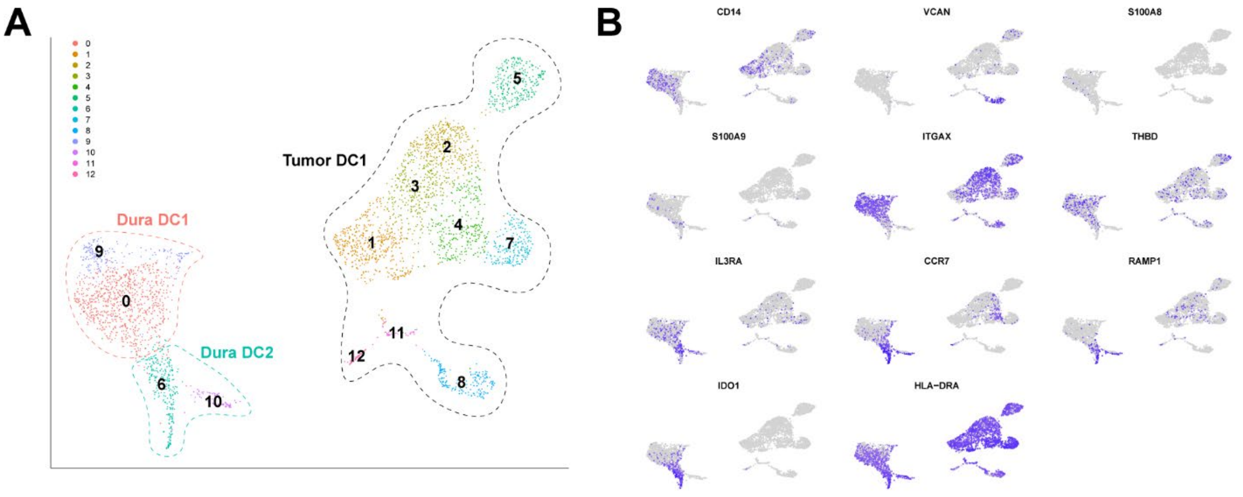


**Fig. S6. Analysis of dura and tumor DCs. (A)** UMAP visualization of dura and tumor DCs. Dura DC1 include C0 and C9. Dura DC2 include C6 and C10. Tumor DC1 include C1, C2, C3, C4, C5, C7, C8, C11, and C12. **(B)** UMAP visualization of dura and tumor DCs for select marker DC genes.

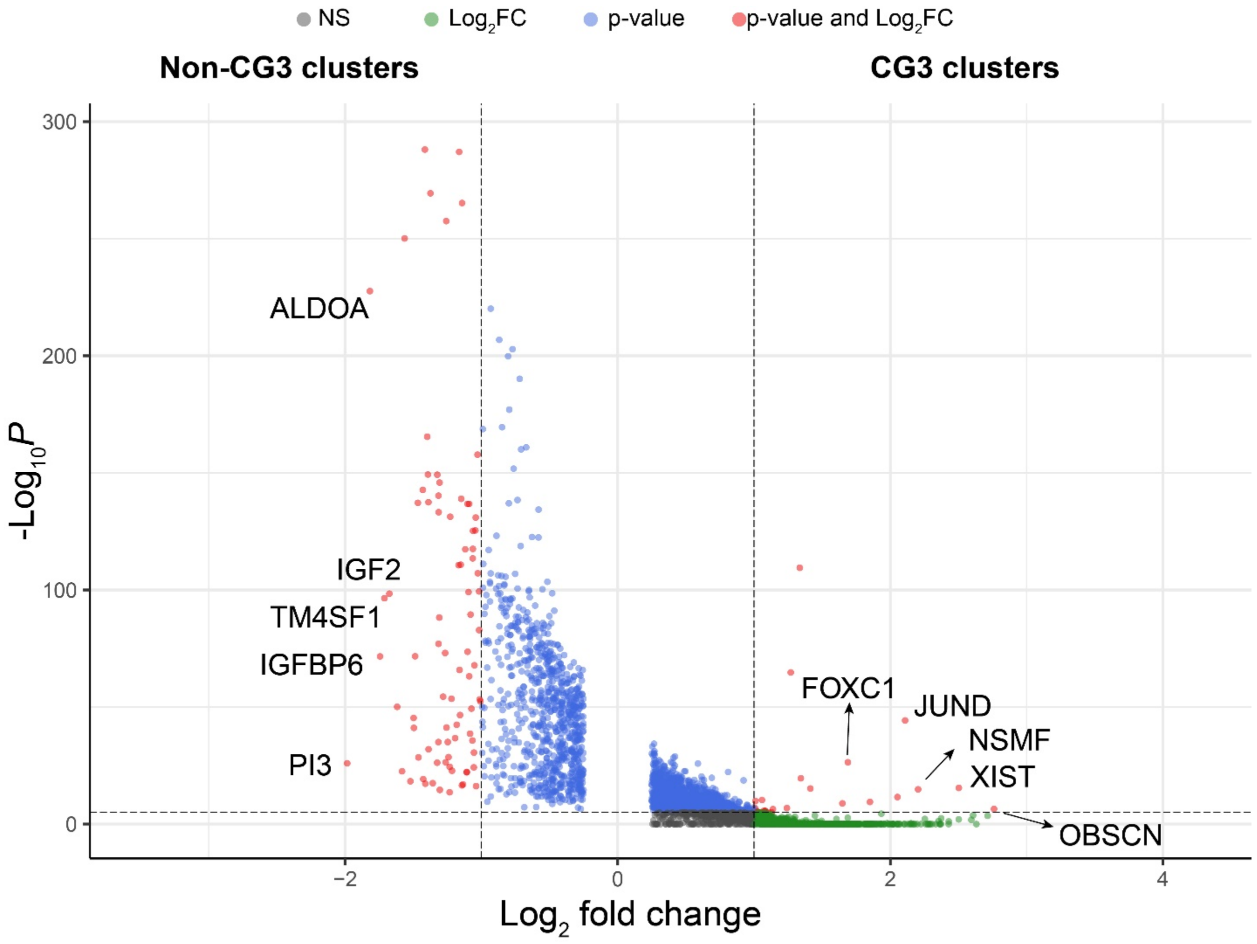


**Fig. S7. Volcano plot of meningioma cluster DEGs.** Volcano plot of genes differentially expressed by CG3 clusters (positive $\log_2$ fold change) and non-CG3 clusters (negative $\log_2$ fold change). $\text{Log}_2$ fold change and p-value thresholds are > |2| and 10e-6, respectively.

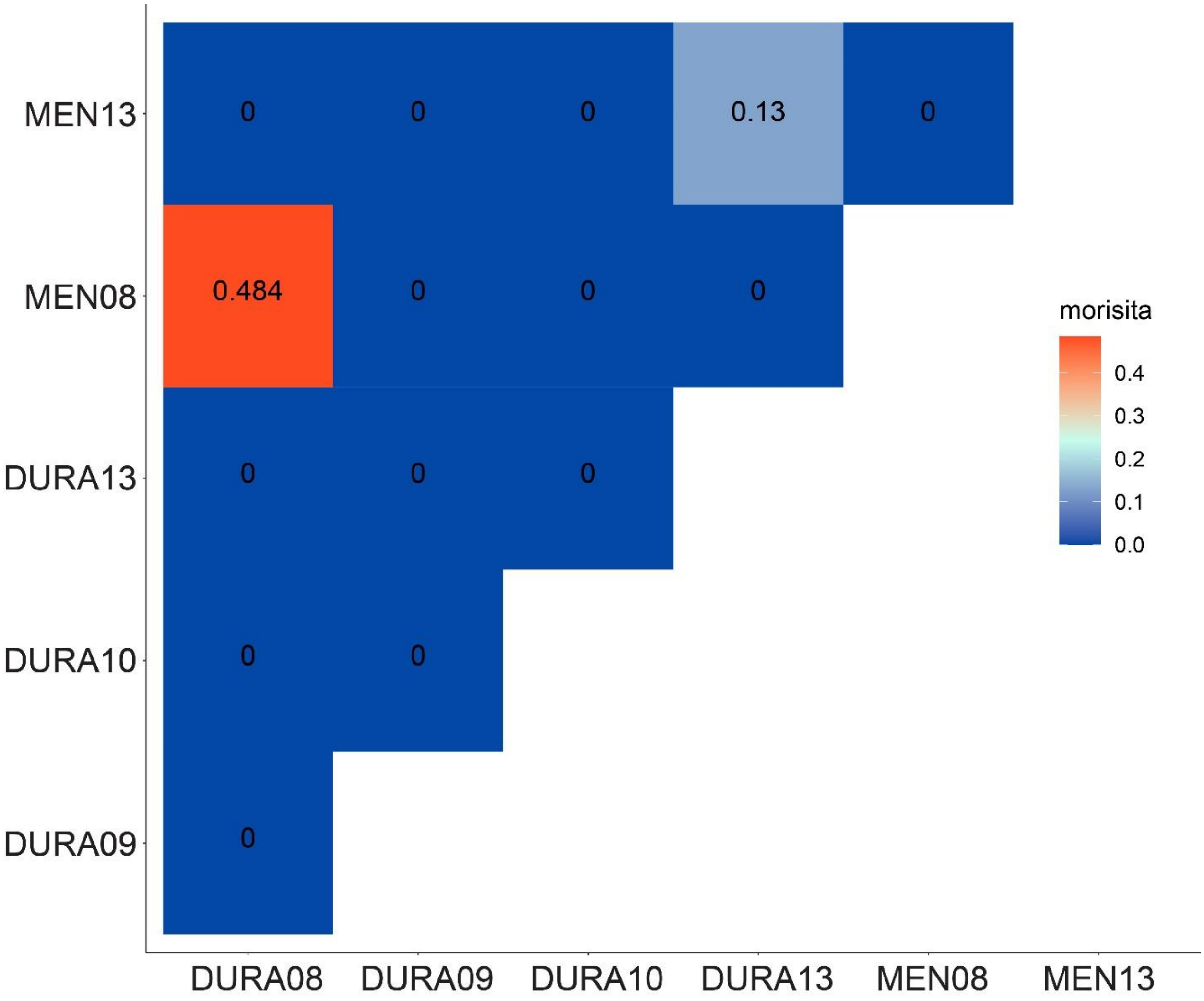


**Fig. S8. Quantification of TCR overlap.** The Morisita overlap index was calculated comparing the overall TCR repertoires from respective dura and tumor samples. Values range from 0 (no similarity) to 1 (completely identical TCR repertoires).

| **Sample Names** | **Patient Gender** | **Tumor Location** | **Pathology** | **Tissue Type** | **Analyses performed** |
|---|---|---|---|---|---|
| **SAMPLE02** | **F** | **L sphenoid wing** | **WHO grade II, atypical** | **Non-tumor associated dura (DURA02)** | **IMC** |
| **SAMPLE05** | **M** | **R sphenoid wing** | **WHO grade II, atypical** | **Non-tumor associated dura (DURA05)** | **3' scRNAseq, IMC** |
| **SAMPLE06** | **F** | **R frontotemporal** | **WHO grade II, atypical** | **Non-tumor associated dura (DURA06)** | **3' scRNAseq** |
| **SAMPLE08** | **M** | **L frontal** | **WHO grade I, angiomatous** | **Non-tumor associated dura (DURA08)** | **5' scRNAseq, VDJ enrichment** |
| | | | | **Meningioma (MEN08)** | **5' scRNAseq, VDJ enrichment** |
| **SAMPLE09** | **F** | **L frontal** | **WHO grade III, anaplastic** | **Non-tumor associated dura (DURA09)** | **5' scRNAseq, VDJ enrichment** |
| | | | | **Meningioma (MEN09)** | **5' scRNAseq** |
| **SAMPLE10** | **M** | **L frontal** | **WHO grade I** | **Non-tumor associated dura (DURA10)** | **5' scRNAseq, VDJ enrichment** |
| | | | | **Meningioma (MEN10)** | **5' scRNAseq** |
| **SAMPLE11** | **M** | **Parafalcine** | **WHO grade II, atypical with patchy angiomatous** | **Non-tumor associated dura (DURA11)** | **5' scRNAseq** |
| | | | | **Meningioma (MEN11)** | **5' scRNAseq** |
| **SAMPLE13** | **F** | **Parafalcine** | **WHO Grade II, atypical** | **Non-tumor associated dura (DURA13)** | **5' scRNAseq, VDJ enrichment** |
| | | | | **Meningioma (MEN13)** | **5' scRNAseq, VDJ enrichment** |

**Table S1. Patient demographics and sample characteristics.** Basic demographic information and sample characteristics.

| Cell type | Markers | Ref. |
|---|---|---|
| Endothelial cell | *PECAM1, CDH5, KDR, SELE, VWF* | 28, 29 |
| Mesenchymal cell | *COL1A1, COL1A2, LUM, DCN, ACTA2, RGS5* | |
| Immune cell | *PTPRC, CD3E, SPI1, CD14* | |
| Naïve/CM T cell | *SELL, CCR7, LEF1, TCF7, KLF2* | 16 |
| CD4+ effector memory T cell | *SELL-, CCR7-, IL7R* | 17 |
| CD8+ effector memory T cell | *SELL-, CCR-, IL7R, CD8A, CD8B* | 17 |
| Resident memory T cell | *CD69, NR4A2, IL7R* | 17 |
| Cytotoxic CD8+ T cell | *PRF1, NKG7, ZNF683, GZMB, CD8A, CD8B* | 16 |
| Natural killer cell | *PRF1, NKG7, GZMB, KLRD1, KLRF1, CD3-* | 18 |
| B cell | *CD79A, MS4A1,* MHC class II | 19 |
| Plasma cell | *IGHG3, IGHA1, DERL1, FKBP11* | 19 |
| Monocyte/macrophage | *CD14, VCAN, S100A8, S100A9, C5AR1, CD68, RNASE1, C1QC, CD163, FCAR, GPNMB,* MHC class II | 20, 21, 22, 23 |
| Myeloid-derived suppressor cell | Monocyte/macrophage genes, MHC class II- | 25 |
| DC1 | *CD14-, ITGAX, THBD,* MHC class II | 20 |
| DC2 | *CD14-, IL3RA, CCR7, IDO1, LAD1, CCL22, RAMP1,* MHC class II | 20, 23 |
| Mast cell | *GATA2, KIT, HPGDS* | 24 |
| Fibroblast | *LUM, DCN, COL1A1, COL1A2, COL3A1* | 28, 29 |
| Mural cell | *ACTA2, MYH11, CNN1, RGS5, PDGFRB, NOTCH3, MCAM, CSPG4* | 29, 30, 31 |

**Table S2. Gene markers.** Markers used to characterize cell identity of obtained clusters based upon current literature.

| Target | Label | Dilution factor | Clone | Manufacturer | Catalog # |
|---|---|---|---|---|---|
| a-SMA | 089Y | 1:50 | 1A4 | eBiosciences | 14-9760-82 |
| CD206 | 141Pr | 1:100 | E2L9N | CST | 91992 |
| CD14 | 144Nd | 1:400 | 3144025D | Fluidigm | EPR3653 |
| CD163 | 147Sm | 1:200 | EDHu-1 | Fluidigm | 3147021D |
| CD11b | 149Sm | 1:100 | EPR1344 | Fluidigm | 3149028D |
| CD31 | 151Eu | 1:75 | EPR3094 | Fluidigm | 3151025D |
| CD4 | 156Gd | 1:450 | EPR6855 | Fluidigm | 3156033D |
| Iba1 | 158Gd | 1:500 | Poly | Novusbio | NBP2-19019 |
| CD68 | 159Tb | 1:550 | KP1 | Fluidigm | 3159035D |
| CD8a | 162Dy | 1:375 | C8/144B | Fluidigm | 3162034D |
| CD45RA | 166Er | 1:1000 | PTPRC/818 | Novusbio | NBP2-47957 |
| GZMB | 167Er | 1:500 | EPR20129-217 | Fluidigm | 3167021D |
| Collagen I | 169Tm | 1:2500 | Polyclonal | Fluidigm | 3169023D |
| CD3 | 170Er | 1:100 | polyclonal C-Terminal | Fluidigm | 3170019D |
| CD45RO | 173Yb | 1:300 | UCHL1 | Fluidigm | 3173016D |
| HLA-DR | 174Yb | 1:350 | LN3 | Fluidigm | 3174025D |
| Histone H3 | 176Yb | 1:2000 | D1H2 | Fluidigm | 3176023D |
| DNA-Ir1 | 191Ir | 1:400 | 125uM | Fluidigm | 201192A |
| DNA-Ir2 | 193Ir | 1:400 | 125uM | Fluidigm | 201192A |

**Table S3. Antibody Information.** Antibody targets with respective conjugated heavy metals, dilution factor, antibody clone, clone number, and manufacturer information. All antibodies were at a concentration of 0.5 mg/mL except for Iba1 (0.336 mg/mL).

| Cell type/polarization state | Markers | Ref. |
|---|---|---|
| Microglia | *AIF1, GPR34, CD14, C1QA, C1QB, C1QC* | 26 |
| Border associated macrophage (BAM) | *MRC1, STAB1, CD163, APOE, MS4A7* | 6 |
| M1 macrophage polarization | *CD86, CD80, IL1R1, TLR2, TLR4, NOS2, SOCS3, IL1B, TNF, IL6, FCGR2A, MARCO, NOS2, NFKB1, STAT1, IRF5, JUN, FCGR1A, IDO1, SOCS1, CXCL10* | 50 |
| M2a macrophage polarization | *CD163, MRC1, TGFB1, SLAMF1, SPHK1, THBS1, HMOX1, STATE3, TLR1, TLR8, TGM2, FCER2, CCL22* | 50 |
| M2b macrophage polarization | *MRC1, TREM2, IGF1, IL1RN, STAT6, KLF2, IRF4, PPARG, PPARD, CD163, TGM2, IL1R2, FCER2, CCL22* | 50 |
| M2c macrophage polarization | *IL6, VEGFA, IGF1, CD86, TNF, FCGR1A, MRC1, TGM2, FCER2, CCL22* | 50 |

**Table S4. Gene markers.** Markers associated with microglia, border associated macrophages, and macrophage polarization states.

**Data S1. DEGs of dura and tumor *CD45*+ and *CD45*- clusters.**

Sheet 1: Results of Wilcoxon rank-sum of each cluster against other clusters present in dura *CD45*+ cells.
Sheet 2: Results of Wilcoxon rank-sum of each cluster against other clusters present in dura *CD45*- cells.
Sheet 3: Results of Wilcoxon rank-sum of each cell type against other cell types present in dura *CD45*- cells.
Sheet 4: Results of Wilcoxon rank-sum of each cluster against other clusters present in pooled dura and tumor *CD45*+ cells.

**Data S2. Analysis of principal component genes of different cell types in dura and tumor samples and associated gene ontology biological pathways.**

Sheet 1: Top 10 genes of the top 10 PCs and associated gene ontology biological pathways of dura DCs.
Sheet 2: Top 15 genes of the top 15 PCs and associated gene ontology biological pathways of dura endothelial cells.
Sheet 3: Top 15 genes of the top 10 PCs and associated gene ontology biological pathways of dura fibroblasts.
Sheet 4: Top 15 genes of the top 15 PCs and associated gene ontology biological pathways of pooled dura and tumor monocyte/macrophages.
Sheet 5: Top 15 genes of the top 15 PCs and associated gene ontology biological pathways of tumor clusters.

**Data S3. Target-ligand interactions output from CellChat.**

List of target-ligand interactions produced from CellChat R package.

**Data S4. Dura and tumor immune cell DEGs.**

Sheet 1: Results of Wilcoxon rank-sum of tumor T cells against dura T cells.
Sheet 2: Results of Wilcoxon rank-sum of tumor NK cells against dura NK cells.

**Data S5. Dura and tumor DC DEGs and meningioma cluster DEGs.**

Sheet 1: Results of Wilcoxon rank-sum of dura DC1 against dura DC2.
Sheet 2: Results of Wilcoxon rank-sum of dura DC1 against tumor DC1.
Sheet 3: Results of Wilcoxon rank-sum of tumor CG2 against tumor not-CG2.
Sheet 4: Results of Wilcoxon rank-sum of tumor CG3 against tumor not-CG3.
